# Optogenetic control of Wnt signaling for modeling early embryogenic patterning with human pluripotent stem cells

**DOI:** 10.1101/665695

**Authors:** Nicole A. Repina, Xiaoping Bao, Joshua A. Zimmermann, David A. Joy, Ravi S. Kane, David V. Schaffer

**Affiliations:** Department of Bioengineering, University of California, Berkeley, California 94720; Graduate Program in Bioengineering, University of California, San Francisco and University of California, Berkeley, California 94720; Department of Chemical and Biomolecular Engineering, University of California, Berkeley, California 94720; Davidson School of Chemical Engineering, Purdue University, West Lafayette, IN 47907; Helen Wills Neuroscience Institute, University of California, Berkeley, California 94720; Department of Molecular and Cell Biology, University of California, Berkeley, California 94720; Gladstone Institute of Cardiovascular Disease, Gladstone Institutes, San Francisco, California 94158; School of Chemical and Biomolecular Engineering, Georgia Institute of Technology, Atlanta, Georgia 30332

## Abstract

The processes of cell proliferation, differentiation, migration, and self-organization during early embryonic development are governed by dynamic, spatially and temporally varying morphogen signals. Analogous tissue patterns emerge spontaneously in embryonic stem cell (ESC) models for gastrulation, but mechanistic insight into this self-organization is limited by a lack of molecular methods to precisely control morphogen signal dynamics. Here we combine optogenetic stimulation and single-cell imaging approaches to study self-organization of human pluripotent stem cells. Precise control of morphogen signal dynamics, achieved through activation of canonical Wnt/β-catenin signaling over a broad high dynamic range (>500-fold) using an optoWnt optogenetic system, drove broad transcriptional changes and mesendoderm differentiation of human ESCs at high efficiency (>95% cells). Furthermore, activating Wnt signaling in subpopulations of ESCs in 2D and 3D cultures induced cell self-organization and morphogenesis reminiscent of human gastrulation, including changes in cell migration and epithelial to mesenchymal transition. Our findings thus reveal an instructive role for Wnt in directing cell patterning in this ESC model for gastrulation.

---

Molecular regulation of embryonic morphogenesis, a process where seemingly identical cells differentiate and organize into spatially defined regions, remains poorly understood in mammalian developmental biology^1^. The emergence of such spatial organization is attributed to cell-intrinsic differences in gene expression^2, 3^ or extrinsic asymmetries within the cell environment^4, 5^. The resulting variability in intracellular signaling leads to changes in cell migration, cell-cell interactions, and/or cell polarity that in turn drive the coordinated organization of specific cell populations^6, 7^.

Self-organization within the embryonic tissue proper, or epiblast, is first evident during gastrulation, where subpopulations of cells reorganize and differentiate along distinct cell lineages to form the three germ layers of the future organism^4^. Initially, asymmetric patterns of molecular signals – morphogens such as Wnt, BMP, Nodal, and FGF – emerge across the morphologically symmetric epiblast of the mouse embryo^4, 8–12^. The establishment of such signaling asymmetry is followed by morphological symmetry-breaking, where a subpopulation of posterior epiblast cells undergoes mesendoderm differentiation and an epithelial to mesenchymal transition (EMT), then migrates away from the epithelial epiblast cells in the region of the primitive streak^13, 14^. During this dynamic process of gastrulation, the temporal order and location of cell migration through the primitive streak is correlated with downstream cell fate outcome, resulting in organization of the developing cell lineages into defined regions within the embryo^13, 15–17^.

However, while advances in live embryo imaging and transcriptomic analysis have unveiled dynamics of cell migration and identified molecular signatures of lineage trajectories^17–19^, a mechanistic and causal understanding of the molecular regulation of gastrulation is lacking. Morphogen signals are necessary for successful gastrulation^8, 10–12^, but which specific signals are sufficient for inducing cell self-organization are unknown. Furthermore, it is unclear how the spatial and temporal dynamics of morphogens regulate cell lineage commitment and link cell fate outcome to migration dynamics through the primitive streak^13, 15–17^.

To gain mechanistic insight into morphogenesis, embryonic stem cell (ESC) culture systems have recently been developed to emulate *in vivo* processes of early mammalian development^20^. Such embryoid or gastruloid models can show remarkable gene expression similarity to natural embryos^21–23^ while enabling molecular perturbation to study mechanism. As one example, aggregates of mouse ESCs (mESCs), derived from pre-implantation epiblast cells^24, 25^, spontaneously self-organize and initiate mesendoderm differentiation, an effect enhanced by an exogenous pulse of Wnt agonist^26–28^. Gastrulation-like events are also observed in mESC aggregates grown adjacent to extraembryonic tissue, an effect suppressed by the Wnt antagonist Dkk1^22, 29^. Excitingly, human ESCs (hESCs) have extended such developmental models from mouse to human embryogenesis, which, due to ethical restrictions, has long been a mystery^20, 30^. For example, hESCs geometrically confined to two-dimensional circular micropatterns self-organize into radially symmetric patterns of germ lineages in response to uniform addition of BMP4 or Wnt agonists,^5, 31, 32^ as well as establish signal feedback loops between Wnt/Nodal/BMP4 pathways^33^. However, as with exogenous molecular perturbation of natural embryos, such ESC-based models rely on uniform addition of signal pathway agonists or inhibitors, which lack dynamic and spatial control. Furthermore, such models are confounded by variations in molecular diffusion of agonists within embryoids^34^, cell variation in morphogen receptor expression^3, 5^, and geometric asymmetries in the cell environment^29, 35^. As a result, mechanistic insight into morphogenesis has been limited in such models since their self-organization is a result of spontaneous and heterogeneous differentiation along diverse cell lineages rather than specific control of cell signaling pathways. How specific morphogen signals direct self-organization, cell fate specification, and migration in gastrulation models thus remains unknown.

To address this critical need for dynamic and specific control of embryonic morphogen signaling, we developed an optogenetic approach to perturb Wnt signaling in hESCs. Light-sensitive protein domains that induce protein-protein interactions or modify protein activity in response to illumination allow optogenetic control of signaling in space and time^36, 37^. Optogenetic strategies have recently been applied for spatiotemporal control of transcription and intracellular signaling in the *Drosophila*^38–41^ 1and zebrafish embryo^42–44^. Here, we implement optogenetic control of canonical Wnt signaling to determine whether differential Wnt signaling can model human gastrulation and lead to emergence of organized shape and structure through collective cell rearrangement. We achieve optogenetic control of Wnt signaling in hESCs by illuminating hESC cultures expressing a fusion of the plant blue-light photoreceptor Cryptochrome 2 (Cry2) to the Wnt co-receptor LRP6^45^. By mimicking differential Wnt presentation with optogenetic stimulation of cell subpopulations, we developed an hESC model for studying Wnt-mediated morphogenesis in the primitive streak. Using this gastrulation model in combination with transcriptomic analysis and single-cell migration studies, we show that Wnt signaling is sufficient for inducing self-organization of cells in an EMT-dependent manner.

## RESULTS

### Optogenetic activation of Wnt/β-catenin signaling in hESCs

Canonical Wnt/β-catenin signaling initiates when extracellular Wnt protein binds to transmembrane receptor Frizzled, an event that triggers multimeric clustering of the Wnt co-receptor low-density lipoprotein receptor-related protein 6 (LRP6)^46^. LRP6 oligomers are subsequently phosphorylated and induce an intracellular signaling cascade that stabilizes the downstream Wnt effector β-catenin, which in turn transcriptionally activates target genes^47^. To render Wnt signaling light-inducible, we previously developed an optogenetic system consisting of the photolyase homology domain of *A. thaliana* blue-light photoreceptor Cryptochrome 2 (Cry2) fused to the cytoplasmic domain of LRP6 (LRP6c)^45^ (Fig. 1a).

**Figure 1.**
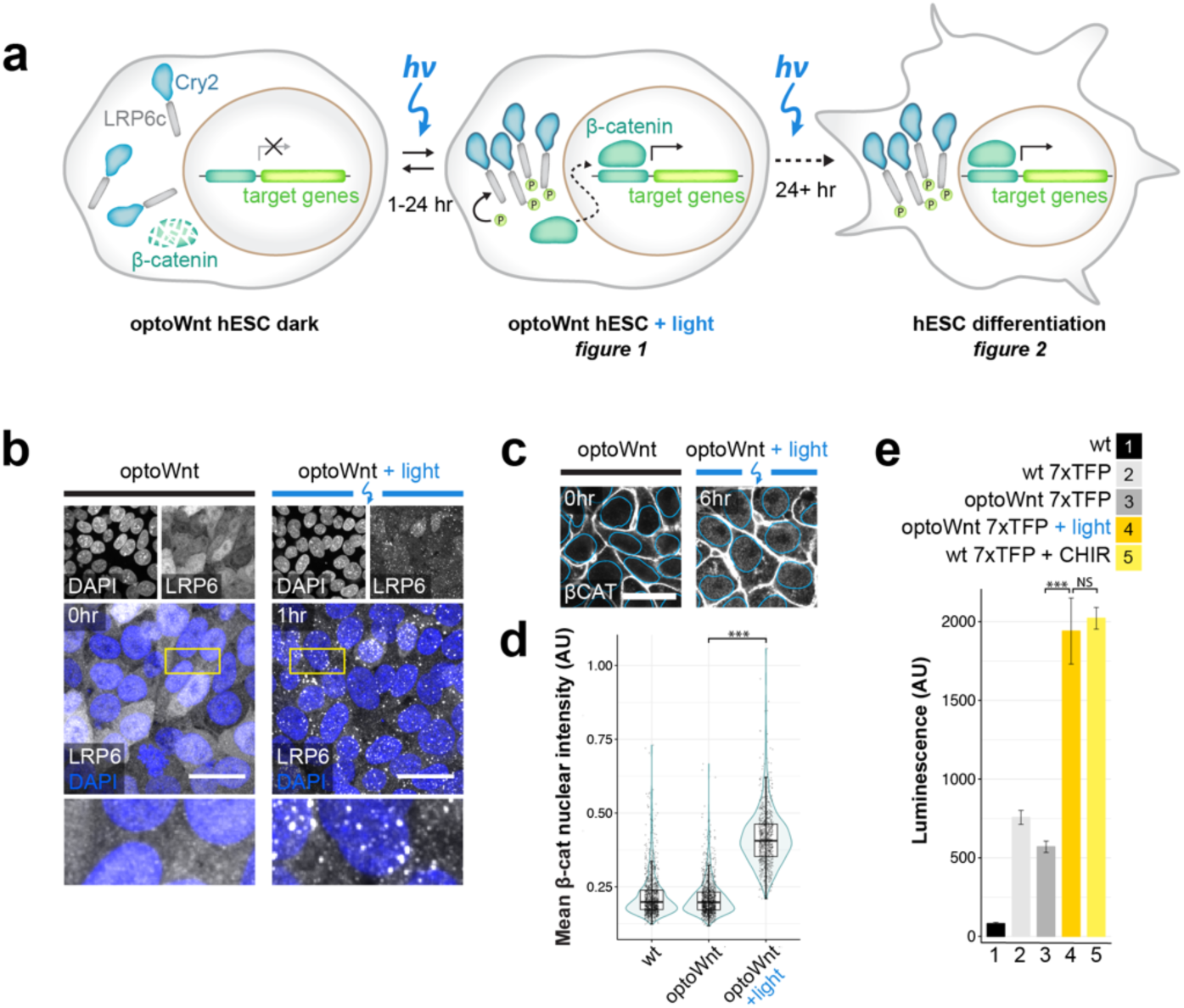
Optogenetic activation of Wnt/β-catenin signaling in hESCs. **a)** Schematic of optogenetic Wnt/β-catenin pathway activation (optoWnt) in hESCs. **b)** Immunostaining for LRP6 in optoWnt hESCs in the dark (left) and after 1 hr of 0.8 *µ*W mm^-2^ illumination (right). Scale bar 25 *µm*. **c)** Representative images of immunostaining for β-catenin in optoWnt hESCs in the dark (left) and after 6 hr illumination (right). Nuclear outline from DAPI stain overlaid in blue. Scale bar 25 *µm*. **d)** Quantification of β-catenin nuclear intensity. Graph shows pooled analysis of 14 fields of view per biological replicate (n = 3), each point represents a single cell. Unpaired two-samples Wilcoxon test (p = 1 x 10^-16^). **e)** Luciferase assay in WT and optoWnt hESCs carrying a 7xTFP reporter for β-catenin activity. Wnt signaling was induced for 24 hr with 0.8 *µ*W mm^-2^ illumination or with CHIR99021 (CHIR, 3µM). ANOVA followed by Tukey test (p = 4.6 x 10^-7^ (3 vs. 4); p = 0.86 (4 vs. 5)). Graph shows mean ± 1 s.d., n = 3 biological replicates.

Here, we extend the application of the Cry2-LRP6c optogenetic system, hereafter named ‘optoWnt’, to hESCs by knocking the optoWnt transgene into the *AAVS1* safe harbor locus for stable expression^48^ (Extended Data Fig. 1a-c). Specifically, we used CRISPR/Cas9-mediated homology-directed repair to generate a clonal, heterozygous hESC line that constitutively expresses Cry2-LRP6c for light-induced Wnt activation and a P2A-linked mCherry for cell identification. The resulting optoWnt hESCs uniformly expressed mCherry and retained a pluripotent phenotype (Extended Data Fig. 1d-e). We also generated clonal induced pluripotent stem cell (iPSC) lines expressing the optoWnt system (Extended Data Fig. 1b-c, f). Blue (470 nm) light stimulation of hESC cultures was achieved with specialized LED illumination devices (light activation at variable amplitude, or LAVA, devices) that allow precise control of the intensity, timing, and uniformity of stimulation (Extended Data Fig. 1g-i).

To confirm that Cry2 is functional in hESCs, we illuminated optoWnt hESC cultures and examined the induction of LRP6 oligomers. In the dark, LRP6 was diffusely distributed throughout the cell cytoplasm, but upon blue light stimulation LRP6 formed distinct oligomers, confirming the functionality of multimeric Cry2 clustering in hESCs (Fig. 1b).

Since activation of canonical Wnt signaling is characterized by β-catenin nuclear translocation in hESCs^47^, we probed for accumulation of nuclear β-catenin upon optogenetic stimulation. From an initial membrane-associated state, reflecting its role in stabilizing adherens junctions, β-catenin protein translocated to the cell cytoplasm and nucleus (Fig. 1c). Single-cell quantification of the mean nuclear intensity of β-catenin showed a significant accumulation of nuclear β-catenin following 6 hrs of optogenetic stimulation (p < 1E-16), with no apparent accumulation in the dark (Fig. 1d).

To assess transcriptional activity of the nuclearly localized β-catenin, we used an established β-catenin-responsive luciferase reporter, 7xTFP^49^, expressed in optoWnt hESCs through lentiviral infection. Consistent with the lack of observable dark-state LRP6 clustering or nuclear β-catenin, unilluminated optoWnt hESCs showed no activation of β-catenin transcriptional activity, suggesting minimal background Wnt pathway activation from optoWnt expression (Fig. 1e). In contrast, illumination over 24 hrs at a constant intensity of 0.8 *µ*W mm^-2^ led to a ∼3-fold increase in luciferase reporter expression (p <1E-7), with activation levels comparable to those induced by the small-molecule Wnt pathway agonist CHIR99021 (CHIR, 3 *µm*).

### OptoWnt stimulation induces hESC differentiation and expression of primitive streak marker Brachyury

Recombinant Wnt3a protein or Wnt pathway small-molecule agonists induce ESC differentiation along a mesendoderm lineage^50, 51^. To determine whether optoWnt can analogously induce hESC and iPSC differentiation, we analyzed the light-induced expression of self-renewal and differentiation markers. We first examined expression of the mesendoderm transcriptional regulator and primitive streak marker Brachyury (BRA, also known by its gene name, *T*), a direct transcriptional target of Wnt signaling^9, 52, 53^. After 48 hrs of light activation, we saw a ∼40-fold increase in BRA/T protein expression in optoWnt hESCs and iPSCs (Fig. 2a, Extended Data Fig. 2a-b). The resulting >99.9% pure BRA+ population was comparable to the CHIR (5 *µm*) positive control, and we did not detect dark-state BRA expression (Fig. 2b).

**Figure 2.**
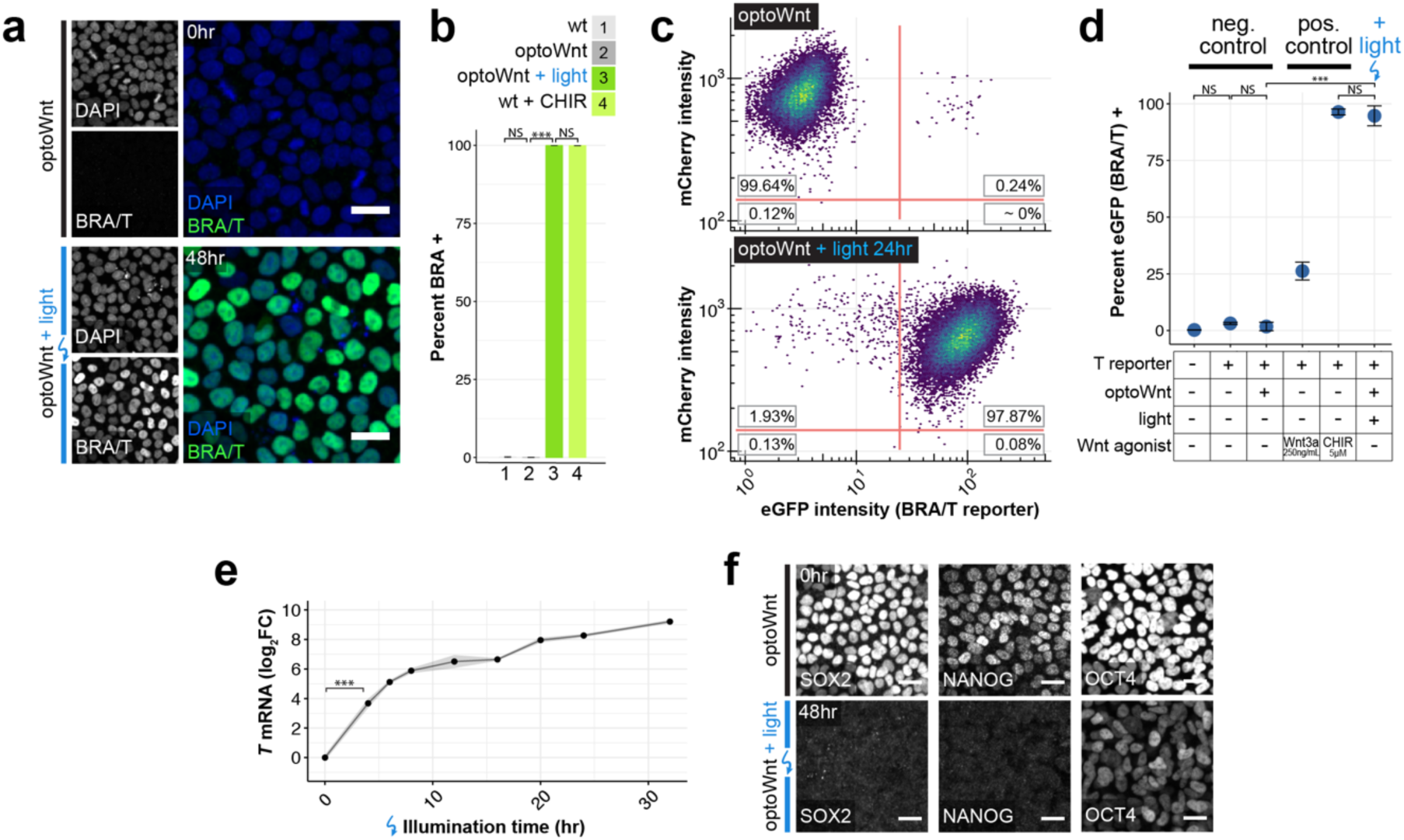
OptoWnt induces BRA expression and hESC differentiation. **a)** Representative images of immunostaining for BRA in optoWnt hESC in the dark (top) or after 48 hrs illumination (bottom). Scale bar 25 *µm*. **b)** Quantification of immunostaining for BRA in WT and optoWnt hESCs following 48 hrs illumination or CHIR (5 *µm*) treatment. Optogenetic stimulation induced >99.9% pure population of BRA+ hESCs. ANOVA followed by Tukey test (p < 1 x 10^-14^ (2 vs. 3); p = 0.28 (1 vs. 2); p = 0.97 (3 vs. 4)). Graph shows mean ± 1 s.d., n = 3 biological replicates. **c)** FACS analysis of optoWnt hESCs (mCherry+) modified with an eGFP reporter at the endogenous BRA/T gene locus (“BRA/T reporter”), kept in the dark (top) or illuminated for 24 hrs (bottom). Graph shows pooled data from 3 biological replicates, ∼30,000 cells per condition. **d)** FACS quantification of percent eGFP-positive cells under indicated conditions for 24 hrs, e.g. Wnt3a recombinant protein (250 ng/mL) or CHIR (5 *µm*). ANOVA followed by Tukey test (p < 10^-12^). Graph shows mean ± 1 s.d., n = 3 to 6 biological replicates. **e)** qPCR timecourse of *T* mRNA expression in optoWnt hESCs at indicated durations of illumination. Graph shows mean log fold change in mRNA expression (log_2_FC) relative to dark (0 hr) condition ± 1 S.E.M, n = 3 biological replicates. **f)** Representative images of immunostaining for pluripotency markers SOX2, NANOG, and OCT4 in optoWnt hESC kept in the dark (top) and following 48 hrs illumination (bottom). Scale bar 25 *µm*.

To better quantify BRA expression at a single-cell level, we generated an hESC reporter cell line co-expressing the optoWnt system and an eGFP reporter for endogenous BRA expression (Extended Data Fig. 1j). Live-cell analysis with flow cytometry showed that > 95% cells expressed BRA after 24 hrs of illumination, with mean eGFP intensity increasing ∼33-fold over unilluminated optoWnt hESCs (Fig. 2c). Quantification of eGFP-positive cells between multiple control conditions showed that signal levels achieved with optoWnt stimulation greatly exceeded activation by high concentrations of recombinant Wnt3a protein (250 ng/mL) and were comparable to 5 *µm* CHIR treatment (Fig. 2d). In contrast, unilluminated optoWnt reporter cells showed no significant eGFP expression relative to wild-type reporter cells, further demonstrating the low dark-state activation. We validated these protein expression results with qPCR for the *T* gene following variable lengths of illumination (Fig. 2e). Onset of *T* transcription was observed after as quickly as 4 hr of illumination (∼14-fold increase in mRNA expression), with expression saturating at ∼500-fold by 32 hr of illumination (Fig. 2e).

Next, we assayed for expression of ESC pluripotency markers SOX2, NANOG, and OCT4. Cells kept in the dark retained pluripotency markers at levels indistinguishable from wild-type (WT) hESCs (Fig. 2f, Extended Data Fig. 2c-f). Conversely, as anticipated for cells undergoing differentiation, optoWnt stimulation led to a decrease in expression of all three pluripotency markers to levels similar to CHIR treatment. The incomplete reduction in OCT4 levels following 48 hrs of illumination is consistent with longer OCT4 persistence during differentiation compared to NANOG or SOX2^54^.

### Global transcriptional profiling of optogenetic stimulation confirms light-induced mesendoderm lineage commitment

To establish a molecular fingerprint for optoWnt-induced differentiation, we measured global transcriptional changes using bulk-population RNA-seq of optoWnt and WT hESCs after 48 hrs of light stimulation (Fig. 3a, **Table S1**). Illuminated WT cells served as a phototoxicity control, and unilluminated optoWnt cells controlled for potential CRISPR/Cas9 knock-in effects, cell perturbation due to optoWnt expression, and dark-state Wnt pathway activation. Principal component analysis (PCA) showed clustering of biological triplicates for each condition and strong transcriptional changes upon optoWnt stimulation that account for 97% gene variance among samples (Fig. 3b). Dark and illuminated WT cells clustered together and differential analysis showed minimal gene expression differences, demonstrating minimal phototoxicity effects after 48 hrs of continuous 0.8 *µ*W mm^-2^ blue light stimulation (Fig 3c). The remaining 3% of variance was captured by the second principle component, which accounted for the slight transcriptional differences between the unilluminated WT vs. optoWnt cells (Fig 3d). Notably, none of the differentially expressed genes (DEGs, highlighted red) are members of the Wnt/β-catenin pathway, suggesting that transcriptional differences are due to gene knock-in or protein overexpression artifacts and not background Wnt pathway activation.

**Figure 3.**
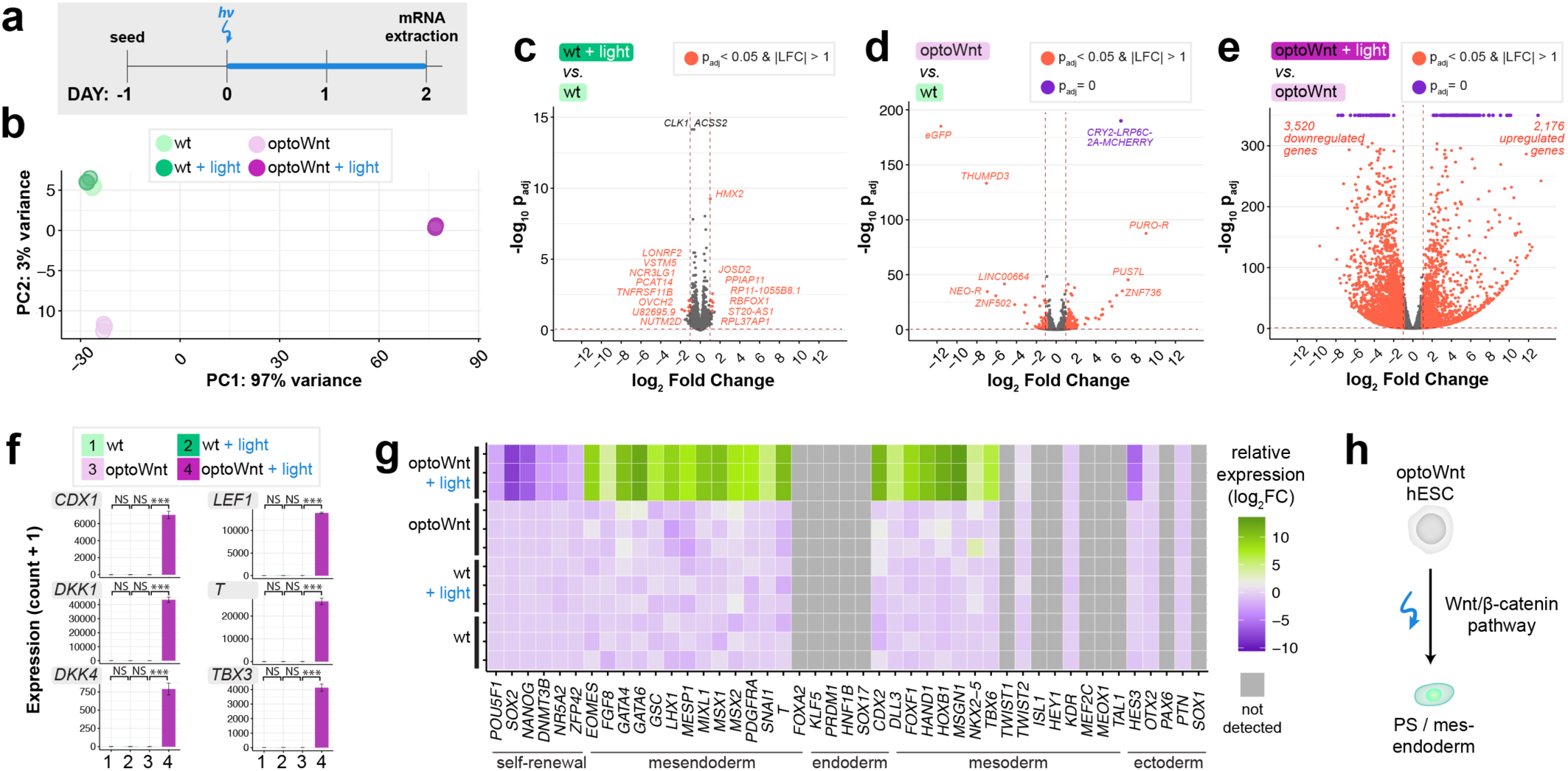
RNA-seq of optoWnt hESCs shows mesendoderm differentiation, low phototoxicity, and low optoWnt dark-state activity. **a)** Schematic of experimental timeline. WT and optoWnt cells were kept in the dark or illuminated at 0.8 *µ*W mm^-2^ for 48 hrs. **b)** PCA of RNA-seq results. Colors designate the four experimental conditions. Each point is a biological replicate. **c-e)** Volcano plots of RNA-seq differential expression analysis, with DEGs (adjusted p-value < 0.05 and log_2_ fold change > 1) highlighted red and DEGs saturated at p = 0 highlighted purple. (c) Illuminated WT vs. unilluminated WT hESCs (phototoxicity control); (d) unilluminated optoWnt vs. unilluminated WT hESCs, (dark-state activity control); (e) illuminated optoWnt vs. unilluminated optoWnt hESCs. **f)** β-catenin target gene expression. ANOVA followed by Tukey test. Graphs show mean expression (read counts + 1) ± 1 s.d., n = 3 biological replicates. **g)** Heat map of log_2_ fold change in lineage markers normalized to WT hESC expression level. Biological replicates displayed for each condition, with undetected genes (read count < 150) shown in grey. **h)** Schematic of light-induced mesendoderm differentiation.

In contrast, optoWnt stimulation induced a broad transcriptional effect, with ∼5,500 differentially expressed genes between the dark and illuminated conditions (Fig 3e). Direct β-catenin target genes, such as *CDX1*, *DKK1*, *T*, and *TBX3*, are among the most differentially expressed genes, all with a log fold change (LFC) of ∼ 9 – 13 (Fig 3f, Extended Data Fig. 3a-b). To determine whether optoWnt stimulation induced hESC differentiation along a mesendoderm lineage, we analyzed transcriptional changes in fate markers associated with embryonic germ layer specification (Fig 3g, Extended Data Fig. 3c-e). OptoWnt stimulation for 48 hrs induced strong upregulation of the primitive streak and mesendoderm markers *T, EOMES, MIXL1*, *GATA6, MSX1, and GATA4* with a corresponding decrease in self-renewal markers *POU5F1, SOX2, and NANOG*. Endodermal markers, such as *FOXA2* and *SOX17*, and ectodermal markers, such as *SOX1* and *PAX6,* remained low or were downregulated. Conversely, certain mesodermal markers, such as *TBX6, FOXF1, and HOXB1*, were upregulated consistent with the role of Wnt in inducing mesoderm differentiation in the absence of high TGFβ signaling^33^. In summary, global RNA sequencing analysis confirmed that optogenetic stimulation of optoWnt hESCs induced robust mesendoderm differentiation with undetectable phototoxicity, low background activity, and large dynamic range (Fig 3h).

### Wnt signaling is sufficient for inducing cell self-organization in 2D and 3D hESC culture

Equipped with a method for optogenetic control of Wnt signaling, we sought to determine whether Wnt signaling is sufficient for inducing cell self-organization. We activated Wnt signaling in a subpopulation of hESCs, allowing us to observe cell interactions between Wnt-active and WT cells. We mixed optoWnt and WT hESCs into a 1:1 heterogeneous culture so that illumination would activate Wnt signaling in only the optoWnt subpopulation of an hESC colony (Fig. 4a). Kept in the dark, the mCherry-positive optoWnt hESCs remained heterogeneously mixed with WT hESCs. Strikingly, illuminated co-cultures showed a strong segregation between the two cell populations, where optoWnt and WT cells separated and created sharp boundaries between aggregates (Fig. 4b, Extended Data Fig. 4a-b). This was a surprising observation because spontaneous Wnt activation has been reported in mESC gastrulation models but did not appear to cause a distinct spatial segregation between cell populations^21, 29^. In addition to segregating, axial cross-sections showed that WT cells remained in an epithelial monolayer while optoWnt cells piled into vertical stacks up to ∼60 *µm* in height and seemed to minimize contact area with WT cells by forming steep boundaries at aggregate edges (Fig. 4b). Illuminated optoWnt cells displayed a mesenchymal morphology, with increased cell protrusions and scattering of single cells from hESC colonies (Extended Data Fig. 4c-d). In contrast, illuminated monocultures of optoWnt cells piled vertically in certain regions with less defined boundaries (Extended Data Fig. 4e). We tested different optoWnt:WT seeding ratios to determine whether there existed an optoWnt dosage threshold for segregation, and we observed clear segregation at all seeding ratios (Extended Data Fig. 5)

**Figure 4.**
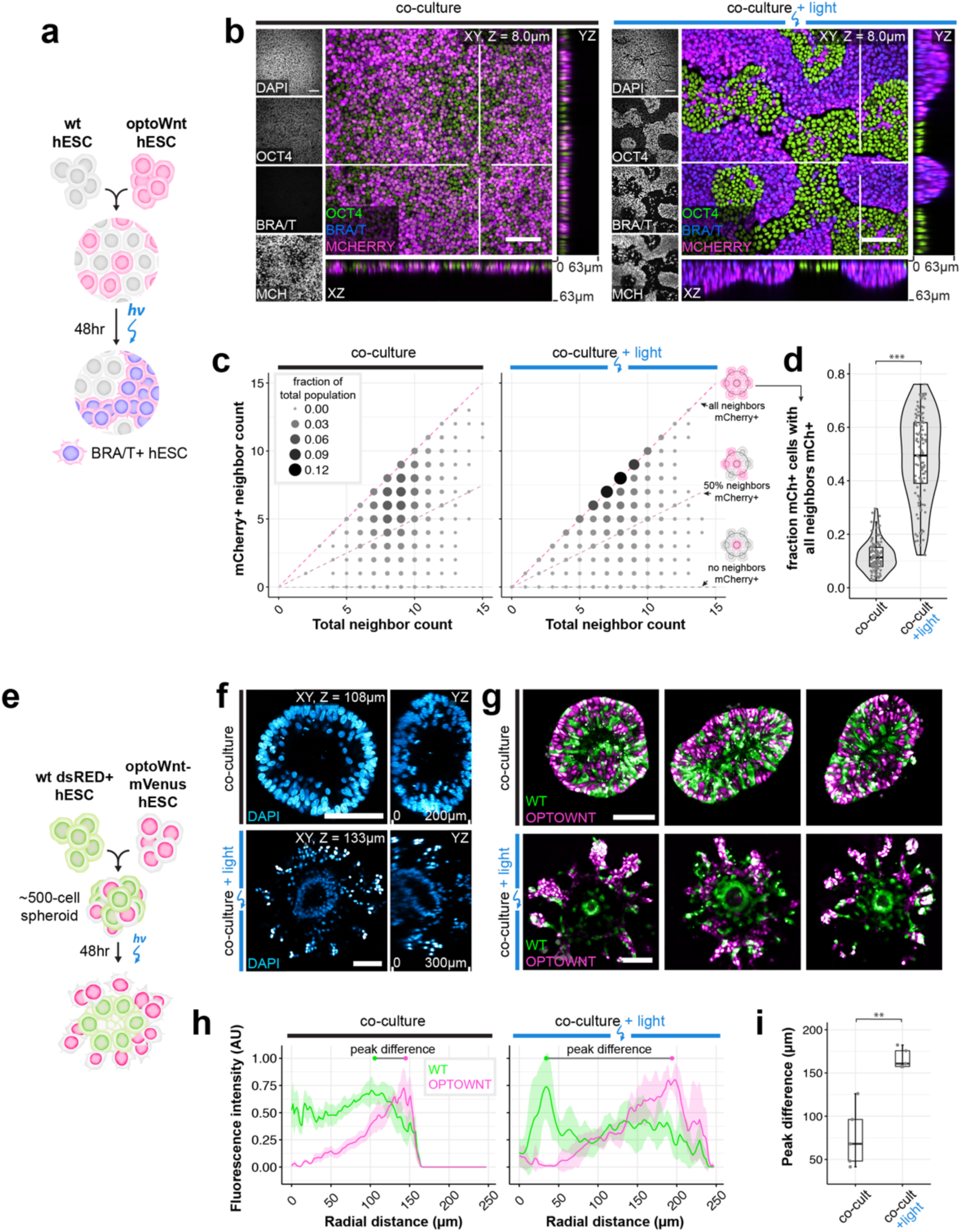
Cell self-organization upon optoWnt stimulation of cell subpopulations. **a)** Schematic of experimental setup of optWnt/WT hESC co-cultures in 2D culture. **b)** Confocal images of optWnt/WT co-cultures in the dark (left panel) and after 48 hrs illumination (right panel), stained for OCT4 and BRA/T. OptoWnt cells are labelled with mCherry (mCh) expression. Scale bar 100 *µm*, YZ and XZ axial cross-sections shown through indicated slices (white lines), 64 *µm* in height. **c)** Cell neighbor analysis of optoWnt (mCh+) cells kept in the dark (left) or illuminated for 48 hrs (right). Graph shows the count of total cell neighbors vs. count of mCh+ cell neighbors across total population of analyzed mCh+ cells (95,685 cells analyzed, pooled analysis from n = 3 biological replicates). Area and color of points is proportional to the fraction of total population.

**Figure 5.**
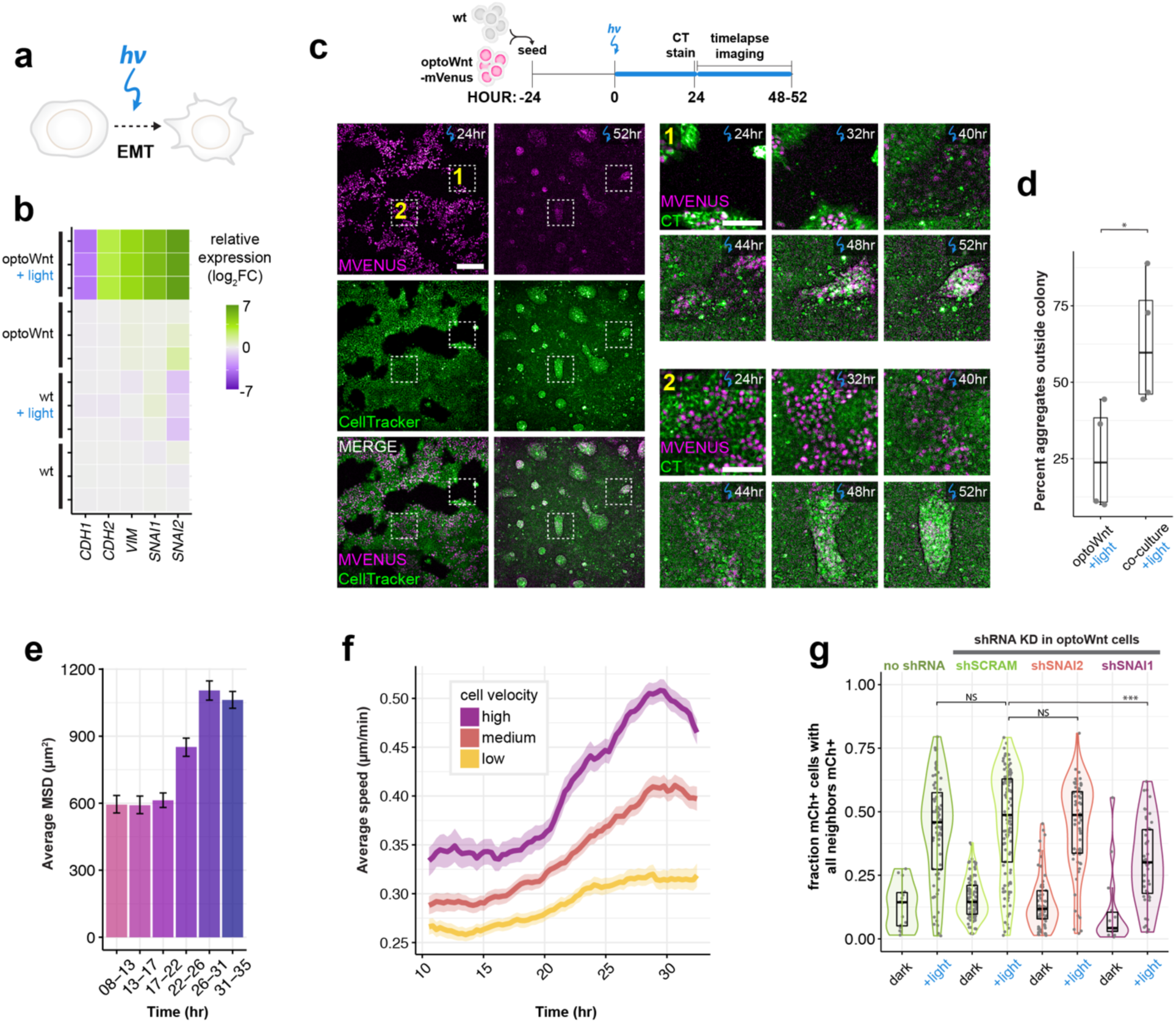
Cell self-organization is mediated by Wnt-induced cell migration and EMT. **a)** Schematic of optogenetic induction of EMT **b)** RNA-seq heat map of log_2_ fold change in EMT marker genes normalized to mean gene expression of WT hESCs. Biological replicates displayed for each condition. **c)** Live-cell timelapse fluorescence imaging of 2D optoWnt/WT co-cultures during optogenetic stimulation at indicated time points. OptoWnt cells are labelled with mVenus-NLS expression, all cells are labelled with CellTracker (CT) dye. Zoom in (right) of regions 1 (cell aggregation outside hESC colony) and 2 (cell aggregation inside hESC colony) at indicated time points. Scale bar 250 *µm* (left), 100 *µm* (zoom in). **d)** Percentage of aggregates forming outside of hESC colonies (as shown in region 1) over total formed aggregates. Graph shows analysis of n = 4 fields of view, ANOVA followed by Tukey test (p = 0.034). **e)** Average mean squared displacement (MSD) of single-cell trajectories of optoWnt cells in optoWnt/WT co-cultures at indicated time intervals after onset of light stimulation. Graph shows mean (>1,000 tracks over 5 fields of view) ± 95% confidence interval. **f)** Average cell speed over time of optoWnt single-cell trajectories binned by median cell velocity (low: 0-0.15 *µm*/min; medium: 0.15-0.18 *µm*/min; high: 0.18-0.6 *µm*/min). Graph shows mean ± 95% confidence interval. **g)** Neighbor analysis of optoWnt/WT co-cultures following shRNA knockdown of EMT regulators *SNAI1* and *SNAI2* in optoWnt cells. Quantification of fraction of optoWnt (mCh+) cells whose neighbors are all mCh+ in co-cultures kept in the dark or illuminated for 48 hrs. Each point represents an analyzed field of view (>16 fields of view analyzed per condition, n = 3 biological replicates). Unpaired two-samples Wilcoxon test (p = 0.00027).

We quantified the extent of segregation by counting the cell neighbors of optoWnt (mCherry+) cells (Fig. 4c, Extended Data Fig. 4f-g). If the two populations were ideally mixed in a 1:1 ratio, the percentage of ‘like’ (i.e. mCherry+) neighbors for each optoWnt cell would average 50%, whereas if the two populations were perfectly segregated, the percentage of ‘like’ neighbors, for optoWnt cells away from population boundaries, would be 100%. Dark co-cultures remained well-mixed with a wide distribution of neighbor counts that were slightly skewed toward higher ‘like’ neighbor percentages, reflecting the clonal expansion that occurs over the 3-day experiment duration (Fig. 4c). In contrast, illuminated co-cultures displayed a strong shift toward optoWnt cells having 100% ‘like’ neighbors, displaying a ∼4-fold increase (p < 0.001) in cells that are entirely surrounded by cells of their own kind, a result consistent with cell segregation (Fig. 4c-d). Strong segregation was observed in all culture media conditions tested, including basal media lacking pluripotency factors like FGF2 and TGFβ, suggesting that Wnt signaling alone is sufficient for driving the observed cell self-organization (Extended Data Fig. 6a-e).

Constant ratios of mCh+ to total neighbors are highlighted with pink and grey lines. **d)** Quantification of fraction of optoWnt (mCh+) cells whose neighbors are all mCh+. Each point represents an analyzed field of view (108 fields of view analyzed, n = 3 biological replicates). Unpaired two-samples Wilcoxon test (p < 2.2 x 10^-16^). **e)** Schematic of experimental setup of optoWnt/WT co-cultures in 3D spheroid cultures. **f)** Two-photon imaging of co-cultures kept in the dark (top) and after 48 hrs illumination (bottom) with DAPI staining of cell nuclei. Scale bars 100 *µm*, YZ axial cross-sections 200 *µm* in height. **g)** Two-photon imaging of co-cultures kept in the dark (top) and after 48 hrs illumination (bottom). WT cells are labelled with dsRed, optoWnt cells are labelled with mVenus-NLS expression. Scale bars 100µm. **h)** Radial quantification of cell segregation in 3D spheroids. Normalized intensity of mean fluorescence signal is graphed as a function of radial distance from spheroid center. Graph shows mean of n = 5 spheroids, ± 1 S.E.M. **i**) Radial distance between peak intensities of WT and optoWnt fluorescence distribution, i.e. ‘peak difference’, labelled in (h). Unpaired two-samples Wilcoxon test (p = 0.0079), n = 5 spheroids.

Self-organization also occurred in three-dimensional (3D) culture when co-cultures of WT and optoWnt hESCs were grown as spheroid aggregates (Fig. 4e). In the dark, WT and optoWnt cells were uniformly distributed throughout the spheroid surrounding a central lumen, as previously observed^55^ (Fig 4f). After illumination, however, spheroids displayed a striking difference in morphology, with a subpopulation of cells that organized radially outward from the central lumen, preserving radial symmetry of the aggregate. Fluorescent labelling of the two cell populations showed that the outer layer of cells was composed of optoWnt cells, while the inner, central lumen was composed of WT cells (Fig 4g**, Supplementary Video 1**). Quantification of the radial distribution of WT (dsRed+) and optoWnt (mVenus-NLS+) cells confirmed segregation of the two populations, as shown by the shift in optoWnt fluorescence toward the spheroid periphery as well as compaction of the WT radial distribution (Fig 4h-i). Further, illuminated optoWnt cells developed cell protrusions that dynamically interacted with the surrounding gel matrix **(Supplementary Video 1**). These results suggest that the lumenized core of WT cells retained an epithelial phenotype, while optoWnt cells obtained a mesenchymal morphology. Such observed self-organization is consistent with our 2D culture results and is reminiscent of the morphogenetic movements of mammalian gastrulation, where mesendoderm similarly segregates radially outward from an epithelial cell layer^56^.

### Self-organization occurs through optogenetic induction of epithelial to mesenchymal transition and cell migration

A key component of embryonic patterning is the coordinated movement of cell populations. Cells of the primitive streak undergo an EMT^14, 57^, a process regulated at least in part by Wnt signaling as mouse embryos mutant in *Wnt3* or β*-catenin* do not ingress or form a primitive streak^8, 58^. To determine whether optoWnt activation is sufficient for EMT induction, we assayed for hallmarks of EMT upon light stimulation (Fig. 5a). Global RNA sequencing showed EMT-associated transcriptional changes upon illumination (Fig. 5b). E-cadherin (*CDH1*) levels markedly decreased after 48hrs of illumination while N-cadherin (*CDH2*) levels increased, a switch in cell adhesion proteins characteristic of cells undergoing EMT^57^. In addition, other indicators of EMT such as cytoskeletal protein Vimentin (*VIM*) and E-cadherin transcriptional repressors Snail and Slug (*SNAI1, SNAI2*) were upregulated. Similar trends were observed at the transcriptional level through qPCR and at the protein level with immunostaining for E- and N-cadherin (Extended Data Fig. 7a-c). These results confirmed that EMT was occurring in response to optoWnt activation.

EMT is often accompanied by a migratory cell phenotype^13, 19, 57^. As clear segregation of cell populations was observed upon optoWnt activation of hESC co-cultures, we sought to analyze the specific role that cell migration during the apparent EMT may have in this self-organization process. We thus performed live-cell timelapse imaging during optogenetic stimulation, in contrast to endpoint analysis, to determine whether cell migration was driving the observed self-organization. Timelapse imaging of 2D cultures showed that optoWnt cells became increasingly migratory and led to visible aggregation after 30-40 hrs of illumination (Fig. 5c**, Supplementary Video 2**). We noticed that the process of segregation occurred through two scenarios, (1) cells migrated out of the epithelial hESC colonies and aggregated in the empty space between colonies or (2) cells migrated and formed aggregates within the colony (Fig. 5d**, Supplementary Video 2-3**). Once formed, the dynamic behavior of aggregates was particularly striking, with aggregates migrating rapidly as a group, merging with nearby aggregates, or detaching from the plate surface entirely (**Supplementary Video 4**). We considered whether apparent segregation could be mediated by an increased proliferation of a rapidly expanding subpopulation of optoWnt cells, but we observed no evidence of clonal expansion in proliferation and imaging studies (Extended Data Fig. 7d-g).

To quantify changes in cell migration upon optoWnt stimulation, we performed single-cell tracking of optoWnt hESCs expressing a nuclearly-localized mVenus (mVenus-NLS) fluorescent tag grown in 2D co-culture with WT hESCs (Extended Data Fig. 8a-c). In the first 24 hrs of illumination, optoWnt hESCs showed no change in cell migration speed, velocity, or mean-squared displacement (MSD), but in the ∼24-32 hr time window these metrics increased ∼2-fold, confirming that cells gain a migratory phenotype upon Wnt stimulation (Fig. 5e, Extended Data Fig. 9a-c). After ∼32 hrs illumination, both MSD and velocity were observed to plateau and then decline in response to aggregation of optoWnt cells, reflecting the increasing role of cellular confinement in determining migration dynamics. Indeed, these time scales align with the emergence of segregation following ∼40 hr of illumination (Fig. 5c, Extended Data Fig. 4d). The ratio of cell displacement over distance and the percentage of time a cell migrated without turning remained constant over time (Extended Data Fig. 9d, i-j), indicating that cells did not increase directionality of their migration. We also observed a large heterogeneity in mean cell migration velocities across the cell population and binned cells into three groups (slow, medium, fast) based on their mean velocity (Extended Data Fig. 9e-h). Over time, the velocity and speed for each group increased but cells maintained their individual migration heterogeneity (Fig. 5f). In summary, these findings support the conclusion that Wnt drives self-organization of differentiating mesendoderm and hESCs through changes in cell migration.

To confirm that self-organization was driven in an EMT-dependent manner, we performed genetic knockdown of target genes in optoWnt hESCs. We were intrigued by the possibility of cell adhesion proteins mediating segregation, a common mechanism for cell sorting ^6, 7^. We found that knockdown of *SNAI1* (encoding Snail protein), a transcriptional repressor of E-cadherin and regulator of EMT, reduced the segregation capacity of optoWnt and WT co-cultures, while knockdown of *SNAI2* (encoding Slug protein) had no detectable effect. (Fig. 5g, Extended Data Fig. 10). Interestingly, these results mimic the phenotype of knockout mice deficient in *Snai1*, which die at gastrulation due to failure to undergo EMT and mesoderm formation^59^, in contrast to *Snai2*-deficient mice, which remain viable^60^. This finding suggests that EMT and E-cadherin downregulation, mediated through *SNAI1*, is required for the observed segregation. In conclusion, we show that differential Wnt signal activation can lead to emergence of organized shape and structure through cell migration and EMT.

## DISCUSSION

Amidst the signaling complexity of the early epiblast and ESC-based models for gastrulation, it has been difficult to tease out the signaling factors responsible for morphogenesis^20^. Wnt signaling induces ESC differentiation and EMT^51, 54, 61^, but whether this can direct the movement of cell subpopulations relative to one another, to generate an organized structure, is unknown. In this study, we combined genetic perturbation, optogenetic stimulation, and live-cell migration tracking to establish a model system for studying early human embryonic patterning. Through optogenetic control of canonical Wnt activity in subpopulations of hESCs, we show that cell-to-cell variation in Wnt signaling is sufficient for inducing cell self-organization and morphogenesis reminiscent of human gastrulation in an EMT-dependent manner.

As biological models become increasingly complex, there is an increased need for tools that can precisely and dynamically perturb a signal input. Previous methods rely on uniform addition of signal pathway agonists or inhibitors, which lack spatial control and are confounded by variations in molecule diffusion and receptor expression^5, 34^. This problem is particularly evident in previous ESC-based organoid or gastruloid models where spontaneous differentiation along heterogeneous cell lineages results in signaling and cell interactions that are poorly controlled^21, 26^. Microfluidics-based approaches can achieve intricate temporal control of morphogen signaling^62^, but such approaches are limiting in their complexity and throughput. In contrast, optogenetics offers the ability to precisely control intracellular signaling pathways in space and time with light. Using engineered methods for optically shaping the illumination pattern, optogenetic control has been recently applied to several developmental systems and achieved spatiotemporal control of ERK signaling in the *Drosophila* embryo^38, 39^ and cell migration in the zebrafish embryo through non-canonical Wnt signaling^42^. Our work extends the optogenetic toolkit to morphogen signaling in mammalian development. Rather than optically shaping the illumination pattern, we introduced subpopulations of optoWnt cells into WT cultures to act as local Wnt signaling hubs upon uniform illumination. By controlling the number or ratio of light-responsive cells in the co-culture, the spatial limits for local signaling density can be investigated. In the future, cell mixing could be combined with cell and light patterning techniques^36, 63^ to achieve intricate geometries and controlled positioning of cells.

Targeting Wnt activation in cell subpopulations allowed us to model several key processes of early embryonic development. First, light stimulation of co-cultures mimicked the presentation of Wnt ligand in the epiblast, which occurs prior to primitive streak formation^9, 64, 65^. From studies of mouse embryogenesis and mESC model systems, Wnt signal induction from a localized source in extraembryonic tissue is thought to break anterior-posterior symmetry of the epiblast^9, 22, 29^, though the source of Wnt signaling remains ambiguous^64, 65^. Alternatively, cell-to-cell variability in the early embryo could also sensitize certain cells to a signal input or elicit spontaneous emergence of collective cell behavior^3, 18, 66^. Our hESC co-cultures model the latter scenario, where individual cells randomly mixed in a population are stimulated with a Wnt signal. That co-cultures are able to self-organize despite their initial salt-and-pepper distribution shows that even in the absence of directional cues, such as gradients of Wnt signaling, heterogeneous activation of Wnt is sufficient for self-organization and mesendoderm migration away from an epithelial cell population (Fig. 4). Consequently, emergence of cell-to-cell heterogeneity in Wnt signaling can lead to tissue organization. This result sheds light on the spontaneous and unexplained self-organization of ESC gastruloid models that occurs in the absence of a localized source of Wnt signaling^21, 26, 27^. Our results suggest that such spontaneous self-organization can emerge from heterogeneous Wnt pathway activation, for example due to variable cell sensitivity to Wnt treatment. Indeed, it has been reported that within a population of ESCs, cells show heterogeneous β-catenin activity^51, 61, 67^ and sensitivity to Wnt agonists^51^, likely a result of inherent gene expression variability or asymmetries in the cell environments. In an embryonic context, epiblast cell heterogeneity is similarly present prior to primitive streak formation^17, 18^ and may thereby help ensure successful morphogenesis in response to a localized Wnt signal.

A second process that our optoWnt/WT co-culture system models is the emergence of a subpopulation of Bra-positive cells adjacent to a subpopulation of Bra-negative cells, mimicking the spatial arrangement of cells at the primitive streak. This model system thus allows studies of the cell-cell interactions and dynamic cell rearrangement between these two populations. Indeed, the directionality of migration in spheroid cultures is consistent with the mouse embryo, where the lumenized epiblast forms a basal lamina on the outer surface of the spheroid through which the mesendoderm cells migrate^56^. However, the absence of a second epithelial layer, equivalent to a hypoblast layer, allows optoWnt cells to continue migrating radially outward instead of becoming spatially confined as in mESC model systems that are bounded by extraembryonic tissues^22^ or in the case of our 2D co-culture model (Fig. 4). Interestingly, since single-cell tracking showed no change in directionality of migration during 2D segregation, the directional migration in 3D cultures is likely a result of the organized epithelial cell polarity of spheroids and/or optoWnt cell interaction with the extracellular matrix. Furthermore, single-cell analysis revealed heterogeneity in optoWnt migration velocity (0.1-0.3µm/min) in response to optoWnt signaling (Extended Data Fig. 15). In the mouse epiblast, cells display similarly heterogeneous migration speeds^13, 19^, with temporal order of migration through the streak and cell displacement correlating with cell fate outcome^13, 15^. Though migration speed heterogeneity in our co-culture models is anticipated due to different local cell densities and confinement in 2D culture, an intriguing developmental analogy could be the spatial patterning of mesoderm fates as cells migrate from the primitive streak to the mesodermal wings.

In conclusion, we establish a framework for studying human gastrulation through optogenetic control of Wnt signaling. As the mechanisms of gastrulation remain poorly understood, this model can be used for further studies of cell-cell interactions and intracellular regulation of cell polarity and migration. We propose that our optogenetic approach can be applied to a variety of stem cell models for mechanistic studies of Wnt signaling dynamics and spatiotemporal signaling thresholds, which would shed light on the ubiquitous Wnt-mediated tissue patterning in developmental and stem cell biology.

## METHODS

### DNA vector assembly

All vectors were constructed using Gibson assembly or standard restriction enzyme cloning. Full sequences and descriptions of constructs used are provided in Supplementary Data. To make all the AAVS1-Pur-CAG-optoWnt plasmids, AAVS1-Pur-CAG-EGFP (Addgene plasmid # 80945) was digested with SalI and MluI and the Cry2-LRP6c construct, containing the photolyase homology region of the *A. thaliana* Cry2 protein^45^, was inserted with standard restriction enzyme cloning. To generate β-catenin luciferase reporter lines, the 7xTFP vector (Addgene plasmid # 24308) was modified to replace puromycin resistance with hygromycin resistance. To generate the Brachyury-2A-eGFP donor plasmid (manuscript under revision), DNA fragments of ∼2 kbp in length were PCR-amplified from the endogenous genomic *T* locus, before and after the stop codon, and were cloned into the OCT4-2A-eGFP donor plasmid (Addgene plasmid #31938). Two sgRNAs targeting at or near *T* stop codon (1: CACTGCATCTTTCGGGACCT<u>GGG</u> and 2: TGGCGACACAGGTGTCCATG<u>AGG</u>) were cloned into the CRISPR-GFP plasmid^68^ via T4 ligation. To generate shRNA knockdown lines, shRNA sequences (Table S3) were subcloned into the pLKO.1 lentiviral expression vector digested with AgeI and EcoRI and modified to express the blastocydin resistance gene and eGFP.

### hESC cell culture

For routine culture and maintenance, all optogenetic and WT hESC lines (H9, WiCell)^30^ and iPSC lines (19-9-7, WiCell)^69^ were grown on Matrigel (Corning, lot # 7268012, 7275006) coated plates in mTeSR1 medium (STEMCELL Technologies) and 1% penicillin/streptomycin (Life Technologies) at 37 °C and 5% CO_2_ with daily media changes. Optogenetic cells were cultured with hood lights off. For illumination experiments, cells were singularized with Accutase (STEMCELL Technologies) at 37 °C for 5 min and seeded onto Matrigel-coated 24-well plates in media containing 5 *µm* ROCK inhibitor Y-27632 (Selleckchem). Cells were seeded at a density of 35k – 70k cell cm^-2^. For co-culture experiments, WT and optoWnt cells were mixed in a 1:1 ratio and seeded at a density of 35k cell cm^-2^. After 20-24 hrs, media was changed to E8 (STEMCELL Technologies), E6 (ThermoFisher), or APEL 2 (STEMCELL Technologies) media without ROCK inhibitor and plates were placed onto LAVA illumination devices and subjected to experimental conditions. When indicated, Wnt agonist CHIR99021 (Stemgent) or recombinant Wnt3a protein (StemRD) was diluted in E8 media and added to cells.

To generate 3D cell spheroids, hESCs (WT H9s and optoWnt cells) were passaged to single cell suspensions with Accutase. A 1:1 ratio of H9s and optoWnt cells in E8 media containing 10 *µm* ROCK inhibitor were added to PDMS inverse-pyramidal microwells to generate 250-cell aggregates^70^. After overnight incubation, aggregates were dislodged from microwells by gentle pipetting. Aggregates were then resuspended in 100% Matrigel and 150 aggregates in 50 *µ*L Matrigel were transferred to 24-well plates. After 10 minutes incubation at 37 °C, pre-warmed E8 was added to each well and exchanged every day for the remainder of the experiment.

### Generation of hESC and iPSC cell lines

Clonal knock-in lines were generated through CRISPR/Cas9-mediated recombination. Prior to nucleofection, hESCs were pre-treated with 10 *µm* ROCK inhibitor for 3 to 4 hours or 5 *µm* Y27632 overnight. Accutase-digested single hESCs were collected and 2.5 - 3.5 million cells were nucleofected with 2.5 *µ*g gRNA AAVS1-T2 (Addgene # 41818), 4.5 *µ*g pCas9-GFP (Addgene #44719), and 6 *µ*g optoWnt donor plasmid in 200 *µ*l room temperature PBS -/- using a Nucleofector^TM^ 2b (Lonza) with program B-016. The resulting cells were plated onto Matrigel-coated 6-well plates containing 3 mL pre-warmed mTeSR1 with 10 *µm* ROCK inhibitor. Once the cells grew to confluency, they were subjected to selection with 1 *µ*g mL^-1^ puromycin in mTeSR1 media for approximately 2 weeks. Clonal lines were generated by picking single-cell clones into wells of a Matrigel-coated 96-well plate that were expanded for 1-2 weeks and subjected to PCR genotyping. As for the Bra-2A-eGFP reporter line, 3 *µ*g T gRNA1, 3 *µ*g T gRNA2 and 6 *µ*g Bra-2A-eGFP donor plasmid were used. After the genotyping, targeted clone was treated with TAT-Cre to remove the PGK-PuroR cassette (manuscript under revision). Dual OptoWnt/Bra reporter lines were generated by CRISPR/Cas9-mediated knock-in of Cry2-LRP6c-2A-mCherry into the *AAVS1* locus of Bra eGFP reporter cells, as described above.

Knock-in at the *AAVS1* locus was verified through PCR on genomic DNA extracted with a Quick-DNA kit (D3024, Zymo Research). For the positive knock-in screen, a band size of 1.1kb was expected using F/R primers 5’ CTGTTTCCCCTTCCCAGGCAGGTCC / 5’ TCGTCGCGGGTGGCGAGGCGCACCG. For determination of zygocity, a band size of 0.2kb was expected for heterozygous clones using F/R primers 5’ CGGTTAATGTGGCTCTGGTT / 5’ GAGAGAGATGGCTCCAGGAA.

The β-catenin luciferase reporter lines were generated through lentiviral infection of WT and optoWnt hESCs with 7x TFP lentivirus packaged and purified from HEK 293T cells. Cells were infected with a multiplicity of infection less than 1 and selected with 50-100 *µ*g mL^-1^ hygromycin for three weeks.

Knock-down lines were generated by lentiviral infection of optoWnt hESCs with shRNA against target genes (Table S3). Infected cells were isolated by FACS sorting for eGFP expression. Knock-down was verified through western blot for target genes.

### Optogenetic stimulation

Cells were seeded on matrigel-coated 24-well plates (0030741021, Eppendorf, black-walled with 170µm coverglass bottom) and placed onto LAVA illumination devices kept in standard 37°C tissue culture incubators. In brief, user-defined illumination patterns were uploaded to the LAVA device for independent illumination control of each well. Unless otherwise noted, optogenetic stimulation was achieved with blue light emitted by arrays of 470nm LEDs continuously illuminating hESCs with 0.8 *µ*W mm^-2^ light for the duration of the experiment (1-48 hrs).

### Immunostaining and imaging

For 2D cell cultures, cells were fixed with 3% paraformaldehyde (ThermoFisher) in PBS for 20min at room temperature and subsequently washed three times with PBS, followed by blocking and permeabilization with 5% donkey serum (D9663, Sigma-Aldrich) and 0.3% Triton X-100 (Fisher Scientific) in PBS (PBS-DT) for 1 hour. Cells were incubated with primary antibodies (Table S2) at 4 °C overnight, then washed three times with PBS, and incubated with fluorescently conjugated secondary antibodies (Invitrogen) at 1:250 dilution for 1 hour at room temperature. Both primary and secondary antibodies were diluted in PBS-DT. Cells were washed with PBS and stained with 0.1 *µ*g mL ^-1^ DAPI nuclear stain (ThermoFisher) prior to imaging. For 3D spheroid cultures, each incubation and wash step was performed overnight at 4 °C overnight to allow for diffusion through Matrigel.

Cell proliferation analysis with EdU was performed by treating cells with 10µM EdU for 20min prior to fixation. Staining for EdU was performed using a Click-iT EdU Alexa Fluor 647 kit (Invitrogen) following manufacturer specifications.

Confocal imaging was performed on a Perkin Elmer Opera Phenix system (QB3 High-Throughput Screening Facility). Brightfield and widefield fluorescence imaging was performed on a Zeiss AxioObserver epi-fluorescent microscope and a Molecular Devices Image Xpress Micro imaging system (CIRM/QB3 Shared Stem Cell Facility). Two-photon imaging of 3D spheroids was performed on a Zeiss LSM 780 AxioExaminer with a Spectra-Physics Mai Tai laser (CRL Molecular Imaging Center).

### Live single-cell imaging and tracking

Co-cultures were treated with CellTracker Red (ThermoFisher) dye diluted 5,000x in mTeSR1 media for 15 min and washed two times. A sealing membrane (Breathe-Easy, Sigma-Aldrich) was applied to plates prior to imaging. Plates were imaged on a Molecular Devices Image Xpress Micro (IXM) imaging system with environmental control (37 °C, 5% CO_2_, and humidity control) using a 10x objective. For each experiment, 4-8 sites were imaged at 10-30 min intervals. For single-cell tracking experiments, cells were imaged at 18 min intervals. Optogenetic stimulation was delivered from the fluorescence light source (SOLA Light Engine, Lumencor) set to 5% intensity, passing through the 10x objective and GFP filter set (472/30nm). Measured power at the sample was 2.82 mW. Optogenetic stimulation was delivered for 3 min at each site prior to imaging of each timepoint (i.e. for 3 min every 10-30 min) in a sequence of short light pulses (500 ms on-pulse, 10 s off-pulse).

Single-cell tracking analysis was performed as follows. To extract cell positions, each the contrast on each image was corrected using adaptive histogram equalization, then the image background was removed using a difference of gaussians filter. Peaks at least 20% above background in the resulting foreground image were then detected using non-local maximum suppression^71^ with a minimum radius of 2.6 *µm*. The cell counts were then fit to an exponential model of cell growth, and frames with anomalous (R^2 > 2500 cells^2) segmentations were discarded. This resulted in a dataset of individual cell detections in 5 fields of view spanning 90 frames between approximately hours 8 and 36 of the stimulation experiment.

Individual cell detections were linked to their nearest neighbor in a radius up to 32.4 *µm* from their previous position both forwards and backwards in time with both a global maximum velocity and neighborhood quasi-rigidity penalty^72^. The resulting track fragments were then iteratively merged with overlapping tracks within 3.4 *µm* and 18 minutes of each other. This process converged after 10 iterations, generating 4,127 total tracks with mean length of 13.5 hours (standard deviation 6.4 hours).

For each track, instantaneous velocity magnitude and direction were approximated using finite differences and then smoothed with a 15 point (4.5 hour) rolling window filter. Cell migration distance was calculated by integrating over finite differences of cell position. Track turning angle was calculated by phase unwrapping change in velocity direction. Finally, periods of persistent migration were determined as those times where a cell was both migrating at least 0.1 *µm*/min and turning no more than 2 degrees/min from its previous velocity direction. Both changes in instantaneous traces and binned values were assessed based on 95% confidence intervals around the mean, as determined by 1000 iterations of bootstrap sampling.

### Image analysis

Microscopy image processing, including stitching and z-slice projection, was performed in Fiji^73^ and image quantification was performed in CellProfiler^74^ with custom analysis pipelines detailed below.

For nuclear detection of β-catenin, nuclei stained with DAPI were identified and used to generate a binary mask applied to the β-catenin image channel. The mean β-catenin intensity per cell nucleus was calculated for each cell in a given field of view. A similar quantification strategy was adopted for quantification of percent positive cells for a given nuclearly-localized cell fate marker (e.g. BRA, SOX2, OCT4, NANOG) with a threshold defining ‘positive’ cells determined from signal intensity of negative controls (see Extended Data Fig. 2).

Quantification of cell segregation in 2D cultures was performed as follows. Nuclear outlines for all cells were identified using the DAPI channel images. Each nucleus was then classified as either mCherry positive or negative based on its mean mCherry nuclear intensity. For each mCherry-positive (mCh+) nucleus, the number of total nuclei and the number of mCh+ nuclei that neighbored the cell nucleus within a 15 *µm* expanded perimeter was calculated using the MeasureObjectNeighbors module in CellProfiler. The distribution of total neighbors vs. mCh+ neighbors was then plotted directly and was also used to calculate the neighbor fraction (i.e. mCh+ neighbors / total neighbors) for each mCh+ nucleus. Next, for each field of view (16-54 per biological replicate), the mCh+ cells whose neighbors were all mCh+ (i.e. neighbor fraction = 1) and who had more than 2 neighbors were counted and normalized to the total number of mCh+ cells in the field of view. This value, fraction mCh+ cells with all neighbors mCh+, was used as a metric for quantifying cell segregation. In a well-mixed population, a small fraction of cells should have a neighbor fraction equal to 1, whereas in a perfectly segregated population, all cells except those located at boundary edges should have a neighbor fraction equal to 1.

Radial quantification of cell distribution in 3D co-culture spheroids was performed in Matlab. First, the center of individual spheroid was calculated. From the center, the pixel intensity of the dsRed or mVenus signal, corresponding to the WT and optoWnt cells, respectively, was determined at 10µm intervals in the radial direction. Locally estimated scatterplot smoothing (Loess) was performed to smooth curves and peak dsRed and mVenus signal was determined by finding the maximum of these curves.

### Luciferase assay

WT and β-catenin reporter hESCs were rinsed with PBS, lysed with lysis buffer (E1531, Promega), and centrifuged to pellet debris. Firefly luciferase expression was quantified by adding 100µL Luciferase Assay Reagent (E1500, Promega) to wells of an opaque 96-well plate containing 20µL lysate supernatant, with resulting luminescence immediately detected on a luminometer (SpectraMax, Molecular Devices). Luminescence intensity was normalized to total protein concentration (Pierce BCA protein assay, Thermo Scientific) to account for proliferation or seeding density variability between samples.

### Flow cytometry and analysis

Cells were lifted with Accutase at 37 °C for 5min, centrifuged, and resuspended in flow buffer (0.5% bovine serum albumin in PBS (w/v)) for analysis. Flow analysis and FACS sorting was performed on a BD Fortessa X20 or BD Aria cell sorter, respectively (CRL Flow Cytometry Facility). Cell counting for proliferation assays was performed using a ThermoFisher Attune (CIRM/QB3 Shared Stem Cell Facility) by measuring the number of cells per unit volume.

Data analysis was performed with FlowJo 10 software. To determine the fraction of BRA-eGFP+ cells after light treatment, gating was set such that less than 0.5% of undifferentiated WT hESCs were positive for eGFP or mCherry.

### RNA extraction, reverse transcription, and qPCR

Cells were lifted with Accutase at 37 °C for 5min, centrifuged, and resuspended in TRI reagent (Zymo Research). To achieve higher RNA yields, two to three wells were pooled, constituting a single biological replicate. RNA was purified using an RNA extraction kit (R2051, Zymo Research) as per manufacturer recommendations with an on-column DNase digestion to remove residual genomic DNA. After measurement of total RNA concentration, 1µg of RNA was converted to cDNA using an iScript cDNA synthesis kit (Bio-Rad). Finally, 10ng of cDNA was used for each SYBR Green qPCR reaction, run in 96-well plate format with a 0.1µM final forward and reverse primer concentration (Table S3). qPCR was conducted for 50 cycles at an annealing temperature of 56 °C on a CFX Connect Real-Time PCR Detection System (Bio-Rad). A melt curve was generated at the end of the PCR reaction and a subset of reactions were run on a 1% agarose gel to ensure that only one product of the expected size was amplified per primer pair. qPCR analysis was conducted by the ddC_t_ method. For each cDNA sample, gene expression was internally normalized to the expression of a human housekeeping gene (RPS18) run on the same qPCR plate. Next, for each gene, expression was normalized to the expression level of WT untreated hESCs. The log of relative expression over this WT control (i.e. log_2_(fold change)) was graphed as a heatmap where color corresponds to mean value of biological replicates. The variability in gene expression was assessed with histogram graphs that show mean and standard deviation of fold change for 3 biological replicates, with at least 2 technical replicates for each biological replicate.

### RNA sequencing and data analysis

RNA was purified as described above, with three biological replicates per experimental condition, and stored at −80 °C until library preparation. Library preparation, sequencing, and data pre-processing were performed by MedGenome Inc (Foster City, CA). In short, libraries were generated with the TruSeq Stranded mRNA Library Prep Kit (Illumina) and sequencing was performed on an Illumina HiSeq 4000 with a sequencing depth of ∼35-65 million aligned reads per sample. Quality of reads was evaluated using the FASTQC tool. Adapter trimming was performed using Fastq-mcf and Cutadapt, followed by contamination removal using Bowtie2. Paired-end reads were aligned to the reference human genome (GRCh38, Ensemble database) using STAR alignment software and raw read counts were estimated using HTSeq. Normalization and differential expression analysis were performed in DESeq2^75^. Genes with an adjusted P-value less than 0.05 and fold change greater than 2 were identified as differentially expressed genes (DEGs). PCA analysis, heatmaps, and volcano plots were generated in R. Unless otherwise stated, fold change in heatmaps was normalized to the WT unilluminated control and calculated using normalized read count + 1 values.

### Western blotting

Cells were lysed in RIPA buffer (Sigma) with phosphatase and protease inhibitors (EMD Millipore). Protein content was measured by bicinchoninic acid (BCA) assay and used to normalize samples to the lowest concentration. Lysates were heated to 95 °C and run on a 4–12% Bis–Tris gel (Life Technologies) and transferred to a nitrocellulose membrane. Membranes were blocked in Odyssey blocking buffer (Li-Cor) and incubated with primary antibody at 4 °C overnight (Table S2). All membrane wash steps were performed using tris-buffered saline with 0.1% Tween-20. Membranes were incubated with secondary antibody, IRDye 800 goat anti-mouse IgG and IRDye 700 goat anti-rabbit IgG (Li-Cor), at a 1:10,000 dilution in blocking buffer for 1 hr at room temperature. Blot fluorescence was visualized using an Odyssey CLx system (Li-Cor) and quantified with the built-in gel analyzer tool in ImageJ.

### Statistical analysis and graphing

Data are presented as mean ± 1 standard deviation (s.d.) unless otherwise specified. Statistical significance was determined by Student’s t-test (two-tail) between two groups, and three or more groups were analyzed by one-way analysis of variance (ANOVA) followed by Tukey test. The unpaired two-samples Wilcoxon test was performed for data that was not normally distributed. P < 0.05 was considered statistically significant (NS P>0.05, *P<0.05, **P<0.01, ***P<0.001). Statistical analysis and data plotting was performed in R.

## ACKNOWLEDGEMENTS

We thank Todd McDevitt and Richard Harland for discussions on stem cell biology and embryology, as well as members of the D.V.S lab and Todd McDevitt lab for helpful discussions. We are grateful to Holly Aaron at the CRL Molecular Imaging Center for microscopy training and assistance, supported by NSF DBI-1041078. We also thank Mary West from the QB3 High-Throughput Screening Facility and CIRM/QB3 Shared Stem Cell Facility and Hector Nolla from the CRL Flow Cytometry Facility for technical assistance. Funding supporting this work was provided by the US National Institutes of Health (to D.V.S), the U.S. National Science Foundation (to N.A.R), the Siebel Scholars Foundation (to N.A.R).

## AUTHOR CONTRIBUTIONS

N.A.R. conceived the study, designed and performed experiments, performed analysis, and wrote the manuscript. X.B. provided intellectual input and performed experiments. J.Z. performed experiments and contributed to analysis. D.J. performed single-cell tracking analysis.

R.S.K conceived the study. D.V.S. conceived the study and wrote the manuscript.

## EXTENDED DATA

This section includes:

- Extended Data Figures 1-10
- Extended Data Table S2-S3
- Description of additional supplementary files (Table S1, Supplementary Videos 1-6)

---

**Extended Data Figure 1.**
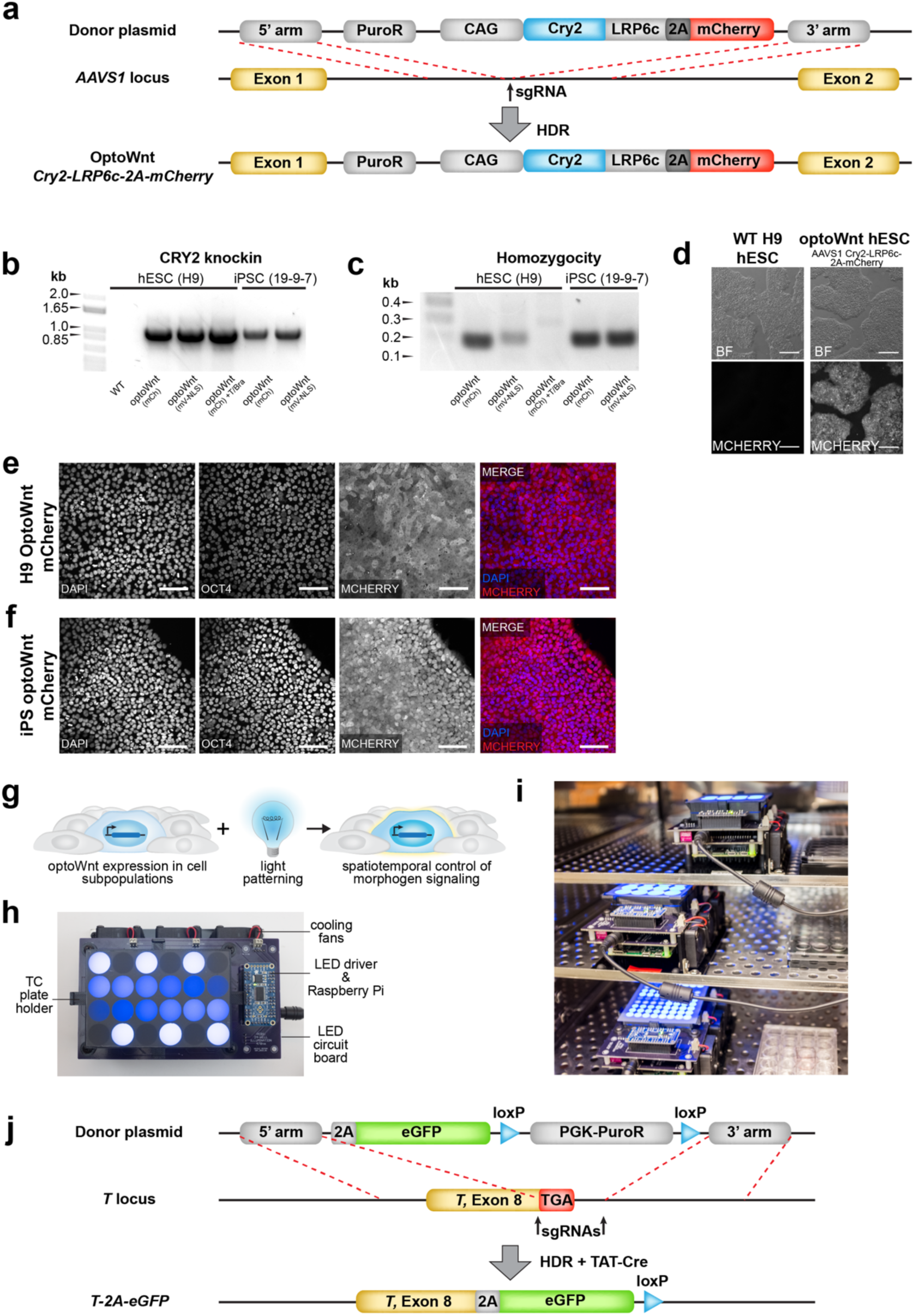
OptoWnt cell line characterization and optical stimulation. **a)** Schematic of optoWnt knock-in strategy using CRISPR-mediated modification of AAVS1 locus. OptoWnt expression driven by synthetic CAG promoter. **b)** PCR genotyping of hESC and iPSC clones after puromycin selection. Expected PCR product for correctly targeted AAVS1 locus is 1.1 kbp. **c)** PCR homozygosity assay on knock-in clones. Clones without a ∼200 bp PCR product were homozygous. **d)** Representative brightfield (BF) and mCherry fluorescence images of live WT and optoWnt hESCs. Scale bar 250 *µm*. **e)** Representative images of immunostaining for OCT4 and mCherry in optoWnt hESCs in routine cell culture, kept in the dark. Scale bar 100 *µm*. **f)** Representative images of immunostaining for OCT4 and mCherry in optoWnt iPSCs in routine cell culture, kept in the dark. Scale bar 100 *µm*. **g)** Schematic of optogenetic experimental setup for stimulation of cell subpopulations. h) Image of illumination device, LAVA board, used for optogenetic stimulation of hESC cultures. Blue light-emitting diodes (LEDs) illuminate a tissue culture (TC) plate placed onto LAVA board. i) Image of LAVA boards kept inside a TC incubator. **j)** Schematic of eGFP knock-in strategy to make BRA/T reporter line using CRISPR-mediated modification of endogenous BRA/T locus.

**Extended Data Figure 2.**
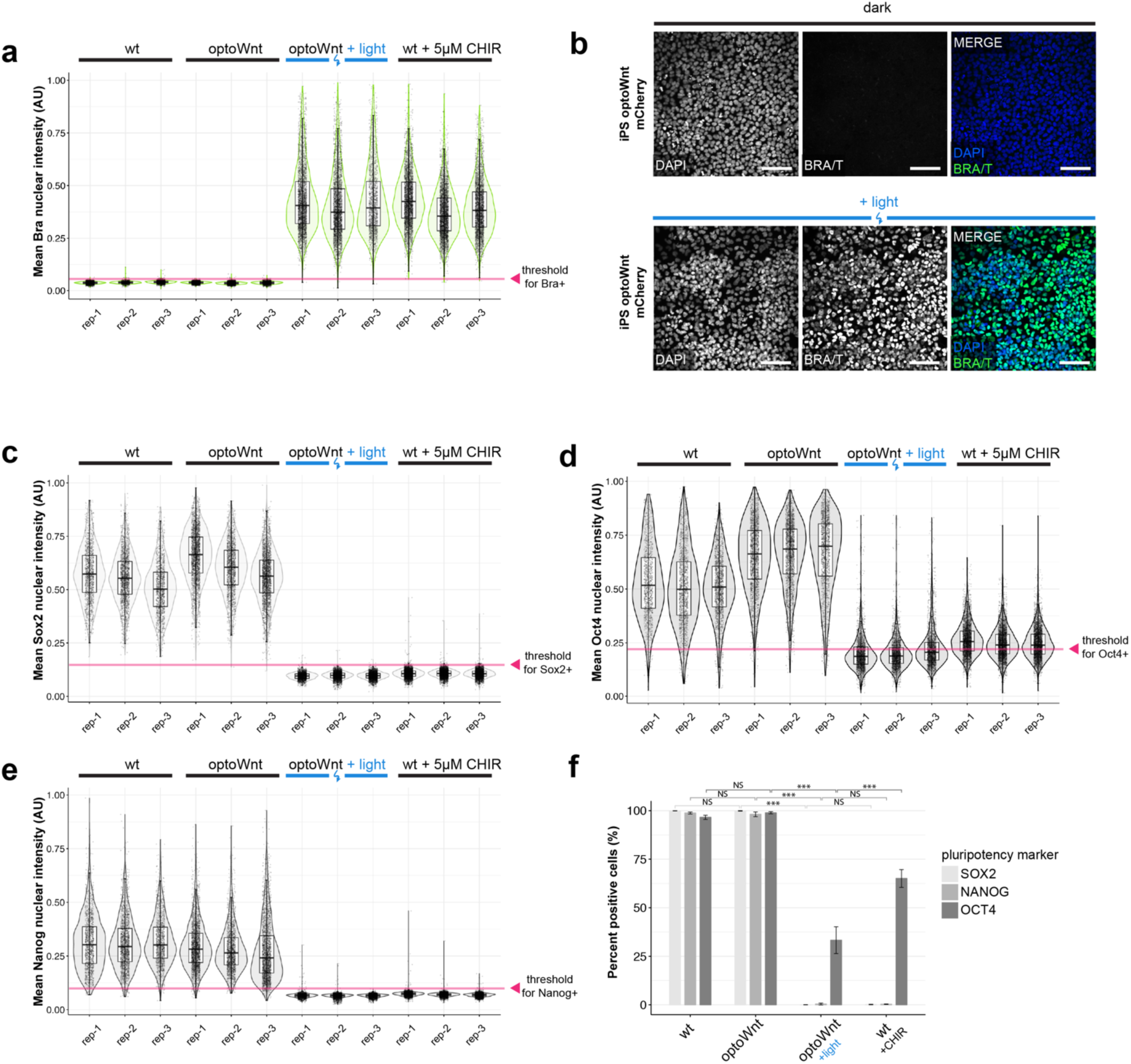
Lineage marker expression in optoWnt hESCs and iPSCs. **a)** Quantification of mean BRA nuclear intensity from immunostaining shown in Fig. 2a-b. Graph shows analysis of biological replicates (rep 1-3) after 48 hrs illumination or CHIR99021 (5 *µm*) treatment. Each point represents a single cell. Threshold for BRA+ classification indicated with red arrow. **b)** Representative images of immunostaining for BRA in optoWnt iPSCs in the dark (top) or after 48 hrs illumination (bottom). Scale bar 100 *µm*. **c-e)** Quantification of mean SOX2 (c), OCT4 (d), and NANOG (e) nuclear intensity from immunostaining shown in Fig. 2f. Graphs show analysis of biological replicates (rep 1-3) after 48 hrs illumination or CHIR99021 (5 *µm*) treatment. Each point represents a single cell. Threshold for SOX2+, OCT4+, or NANOG+ classification indicated with red arrow. **f)** Quantification of percent positive cells for pluripotency markers based on immunostaining shown in (c)-(e) and Fig. 2f. Graph shows percent positive cells ± 1 s.d., n = 3 biological replicates. ANOVA followed by Tukey test.

**Extended Data Figure 3.**
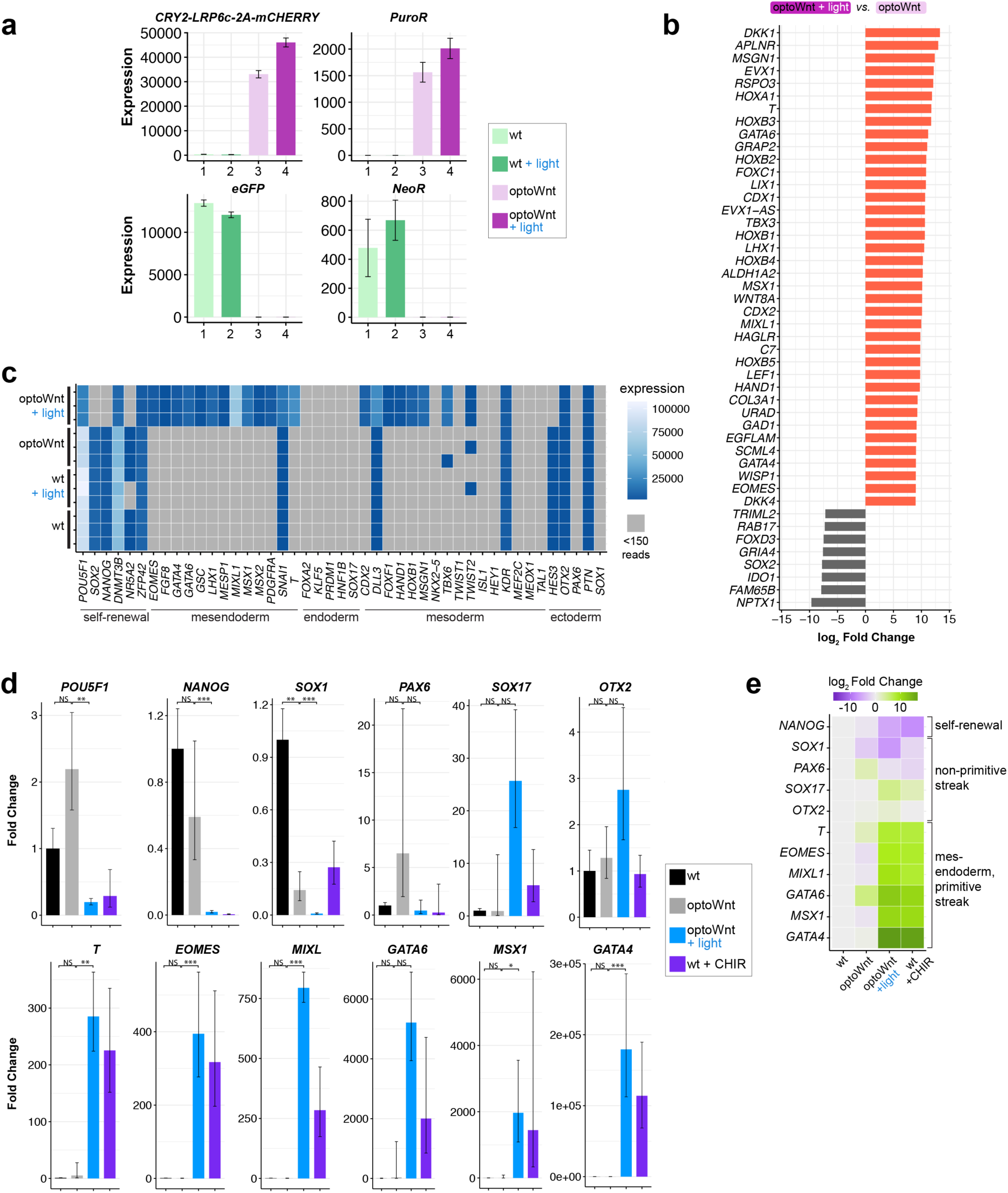
Validation of RNA-seq study of optoWnt-induced hESC differentiation. **a)** RNA-seq results of indicated genes that mark WT (eGFP+, Neomycin-resistant) and optoWnt (Cry-LRP6c-2A-mCherry+, Puromycin-resistant) cells. Graphs show mean expression (read count + 1) ± 1 s.d., n = 3 biological replicates. **b)** Top upregulated (red) and downregulated (grey) genes in illuminated optoWnt vs. unilluminated optoWnt hESCs. Graph shows mean log_2_ fold change for each indicated gene. **c)** Heat map of mRNA expression (read count + 1) of indicated lineage markers. Biological replicates displayed for each condition, with undetected genes (read count < 150) shown in grey. **d)** qPCR validation of RNA-seq results with indicated lineage markers and comparison to CHIR (3µM) treatment. Graphs show mean fold change ± 1 s.d., n = 3 biological replicates. ANOVA followed by Tukey test. **e)** Heat map of mean log2 fold change in lineage markers normalized to WT expression level, from qPCR data shown in (d).

**Extended Data Figure 4.**
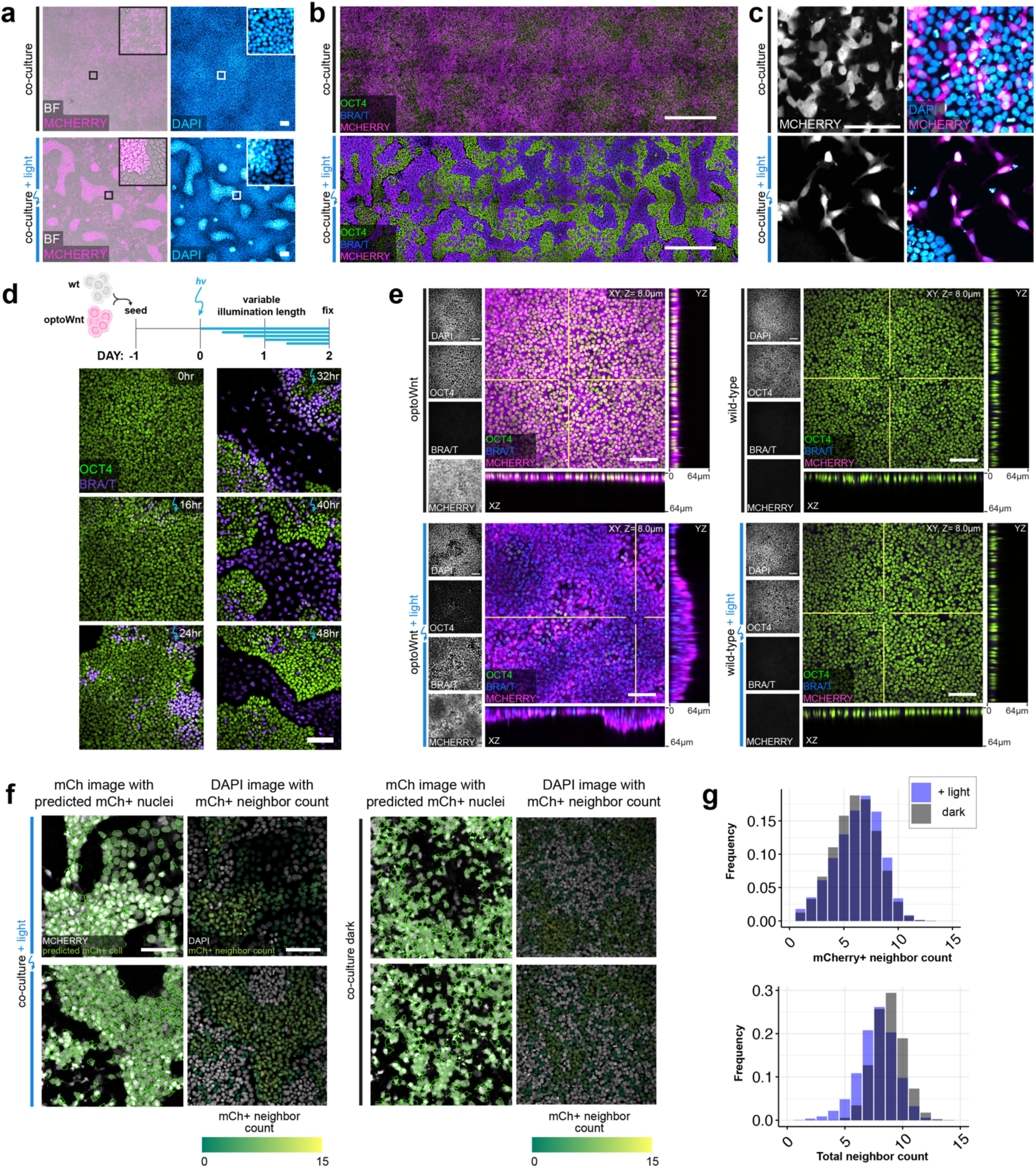
Quantification of cell self-organization upon optoWnt stimulation of cell subpopulations. **a)** Representative images of brightfield (BF), mCherry fluorescence (expressed in optoWnt cells), and DAPI nuclear stain of fixed optoWnt/WT co-cultures in the dark (top panel) after 48 hrs of illumination (bottom panel). Scale bar 100 *µm*. **b)** Stitched images of optoWnt/WT co-cultures show large-scale pattern of cell self-organization. Scale bar 500 *µm*. **c)** Representative image of optoWnt/WT co-cultures seeded at lower cell density (20k cell cm^-2^) show single-cell scattering and morphology changes in optoWnt (mCh+) cells after 48 hrs illumination. Scale bar 100 *µm*. **d)** Representative fluorescence images of optoWnt/WT co-cultures after indicated durations of illumination, stained for OCT and BRA/T. Scale bar 100 *µm*. **e)** Confocal images of optoWnt monocultures (left panels) and WT monocultures (right panels) in the dark or after 48 hrs illumination, stained for OCT4 and BRA/T. OptoWnt cells labelled with mCh. Scale bar 100 *µm*, YZ and XZ axial cross-sections shown through indicated slices (white lines), 64 *µm* in height. **f)** Sample images of cell neighbor analysis in CellProfiler of illuminated (left panel) and dark (right panel) co-cultures. Nuclear outline of mCh+ nuclei (green) overlaid on mCh channel image (left column). DAPI channel image (right column) shown with overlay of mCh+ nuclei colored by mCh+ positive neighbor count. Scale bar 100 *µm*. **g)** Histogram of total cell neighbor counts (bottom) and mCh+ cell neighbor counts (top) across all analyzed cells show comparable cell densities between light and dark conditions.

**Extended Data Figure 5.**
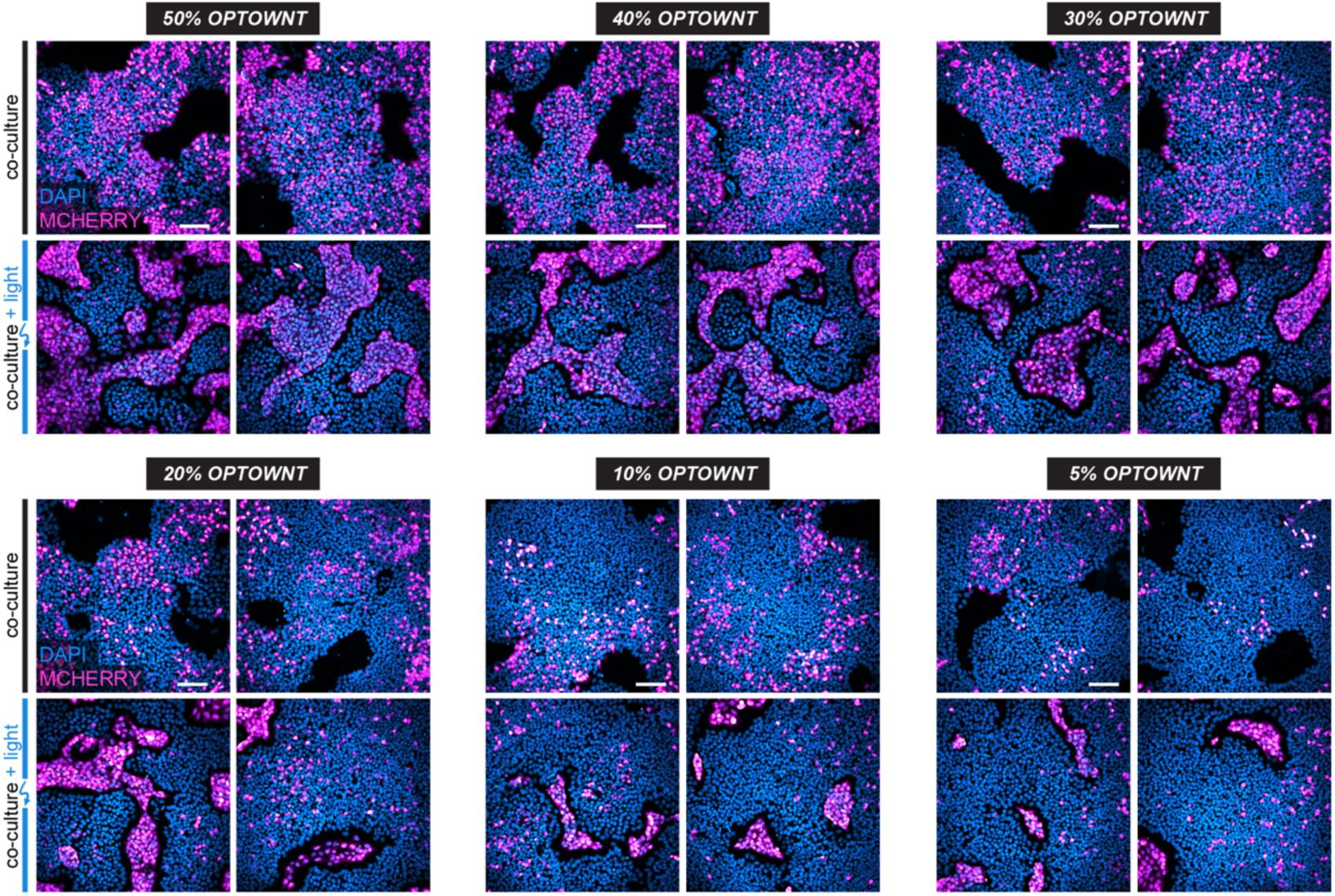
Cell self-organization evident at variable dosages of optoWnt cells in optoWnt/WT co-cultures. Representative fluorescence images of optoWnt/WT co-cultures at indicated seeding doses (e.g. 40% optoWnt indicates 2:3 ratio of optoWnt:WT cells). OptoWnt cells are mCh+. Scale bar 100 *µm*.

**Extended Data Figure 6.**
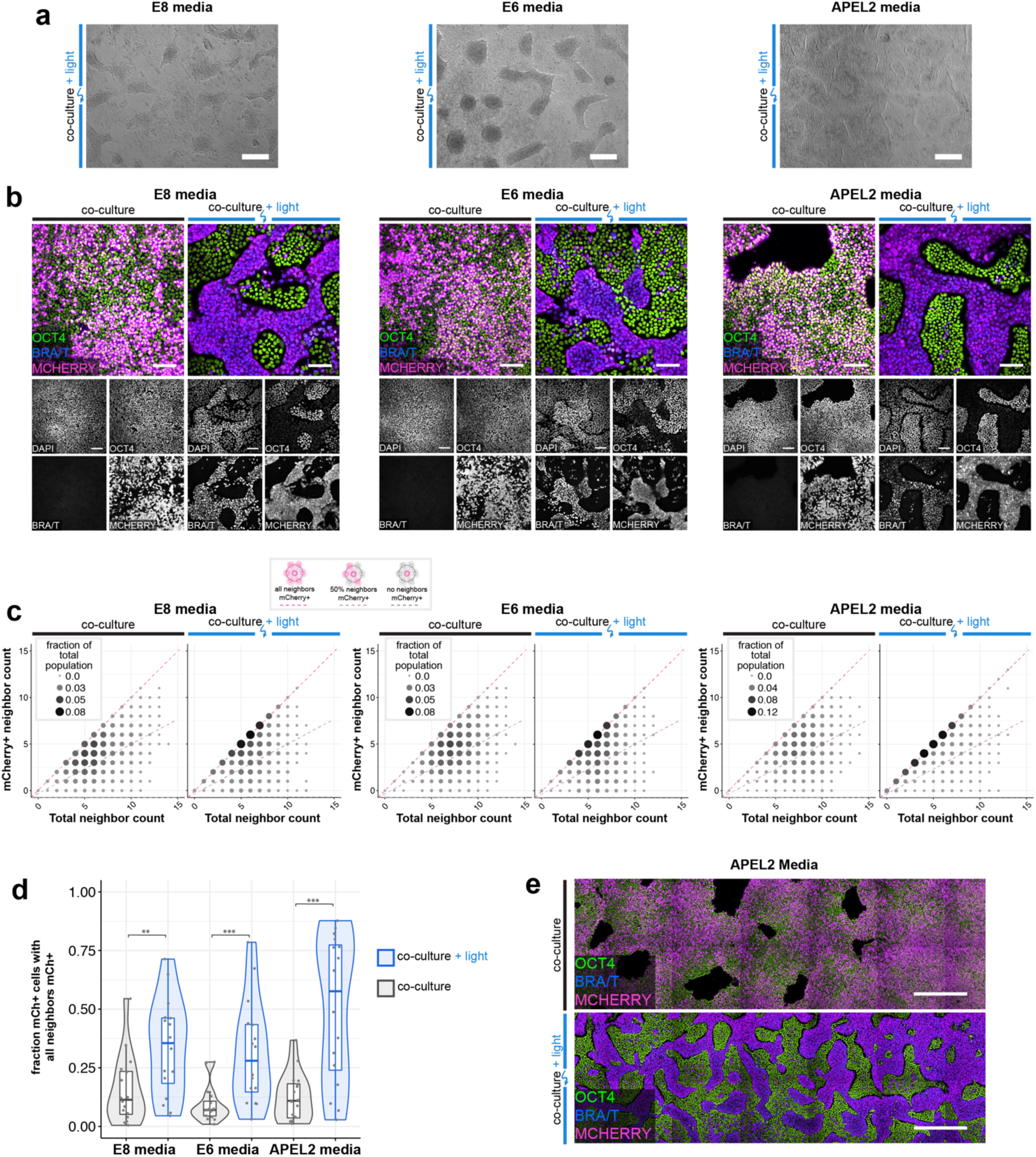
Cell self-organization occurs in media without FGF and TGFβ agonists. **a)** Representative brightfield images of optoWnt/WT co-cultures after 48 hrs illumination in E8, E6, and APEL2 media. Scale bar 250 *µm*. **b)** Confocal images of optoWnt/WT co-cultures in indicated media conditions, stained for OCT4 and BRA/T. OptoWnt cells labelled with mCh. Scale bar 100 *µm*. **c)** Cell neighbor analysis of optoWnt (mCh+) cells in co-culture. Graph shows count of total cell neighbors vs. count of mCh+ cell neighbors across total population of analyzed mCh+ cells (25,742 cells analyzed, pooled analysis from n=3 biological replicates for each condition). Area and color of points is proportional to the fraction of total population. Constant ratios of mCh+ to total neighbors are highlighted with pink and grey lines. **d)** Quantification of fraction of optoWnt (mCh+) cells whose neighbors are all mCh+. Each point represents an analyzed field of view (16 fields of view analyzed per condition, n=3 biological replicates). Unpaired two-samples Wilcoxon test (p_E8_ = 0.011; p_E6_ = 3.3 x 10^-4^; p_APEL2_ = 3.9 x 10^-4^). **e)** Stitched images of optoWnt/WT co-cultures in APEL2 media show large-scale pattern of cell self-organization. Scale bar 500 *µm*.

**Extended Data Figure 7.**
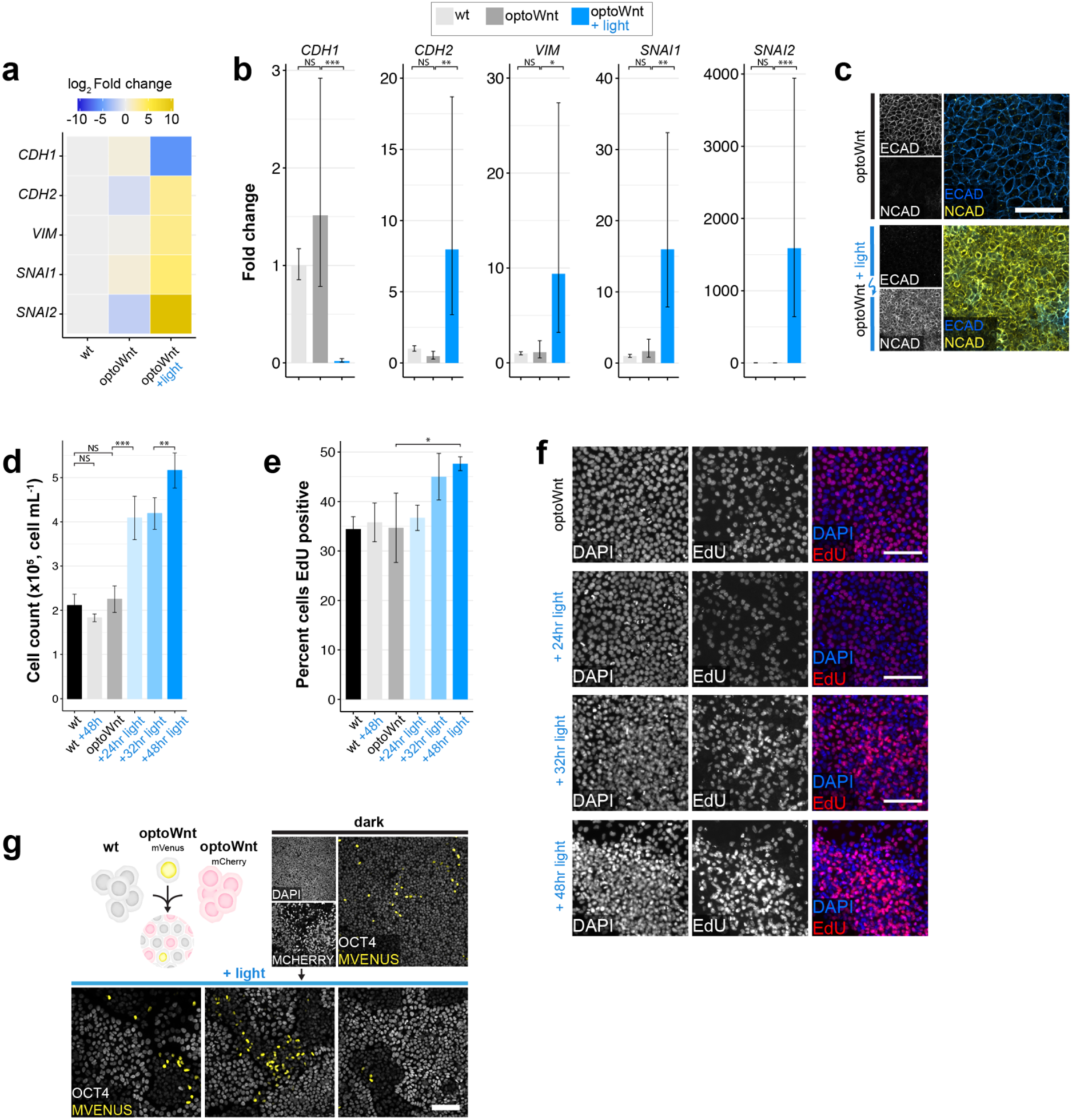
EMT and increased cell proliferation upon optoWnt stimulation. **a-b)** qPCR for EMT markers in WT and optoWnt hESC monocultures cultured in APEL2 media. Graphs show heatmap of mean log2 fold change (a) and mean fold change (b) over WT hESCs ± 1 s.d., n = 3 biological replicates. ANOVA followed by Tukey test. **c)** Representative images of immunostaining for ECAD and NCAD in unilluminated (top) and illuminated (bottom) optoWnt cells. Scale bar 100 *µm*. **d)** Endpoint cell count of optoWnt cells after indicated illumination duration. Graph shows mean ± 1 s.d., n=3 replicates. ANOVA followed by Tukey test. **e-f)** EdU stain of optoWnt cells after indicated illumination duration, with quantification of percent EdU+ cells (e) and representative images of EdU staining (f). Graph shows mean ± 1 s.d., n=3 replicates. ANOVA followed by Tukey test. Scale bar 100 *µm*. **g)** No observed clonal expansion of optoWnt cells when optoWnt-mVenus-NLS cells were dosed into optoWnt-mCh/WT co-cultures. Scale bar 100µm.

**Extended Data Figure 8.**
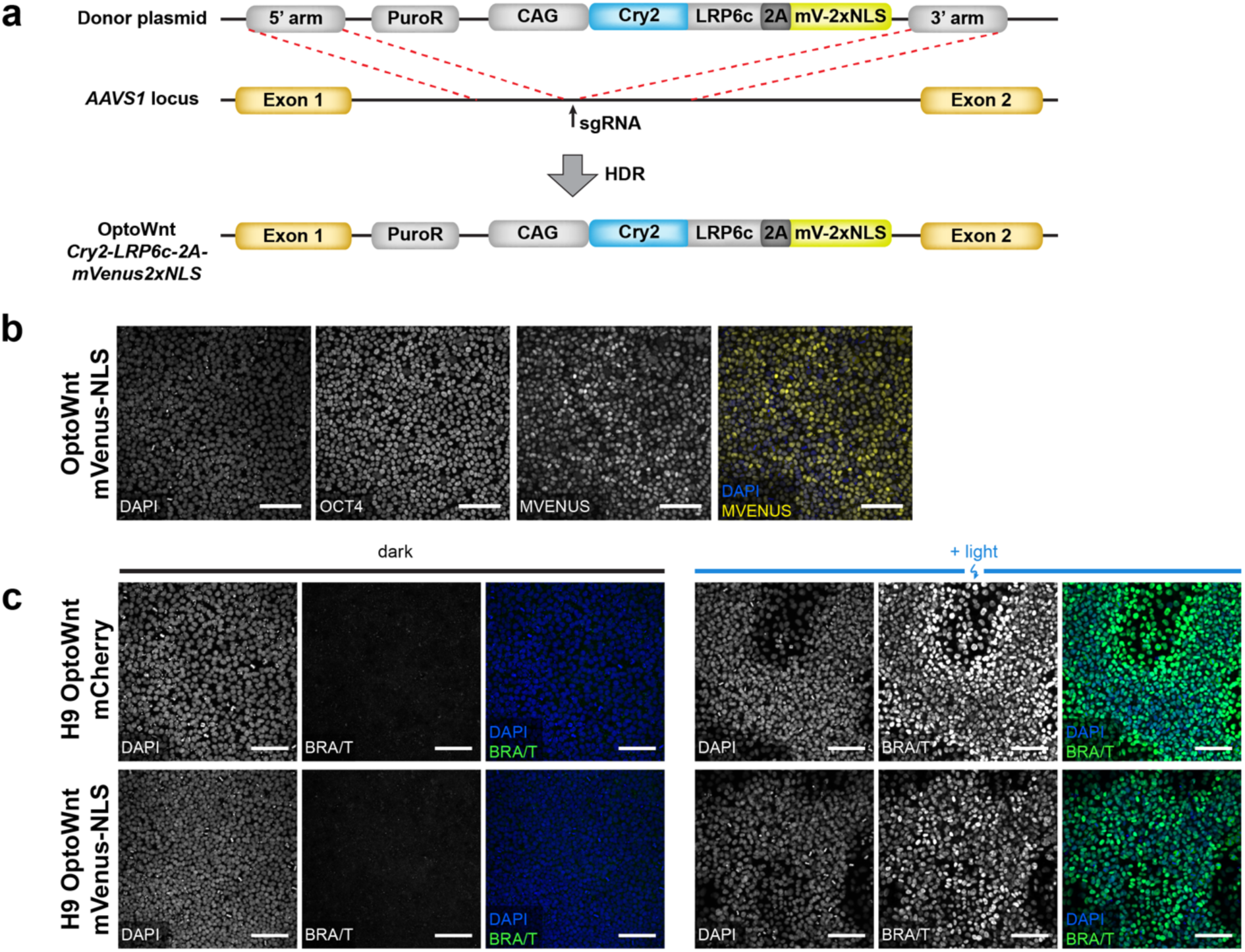
Characterization of optoWnt-mVenus-NLS hESC line. **a)** Schematic diagram of knock-in strategy at the AAVS1 locus. **b)** Representative images of immunostaining for OCT4 and mVenus in optoWnt-mVenus-NLS hESCs in routine cell culture, kept in the dark. Scale bar 100 *µm*. **c)** Light-induced BRA expression of optoWnt-mVenus-NLS line is comparable to optoWnt-mCherry line. Scale bars 100µm.

**Extended Data Figure 9.**
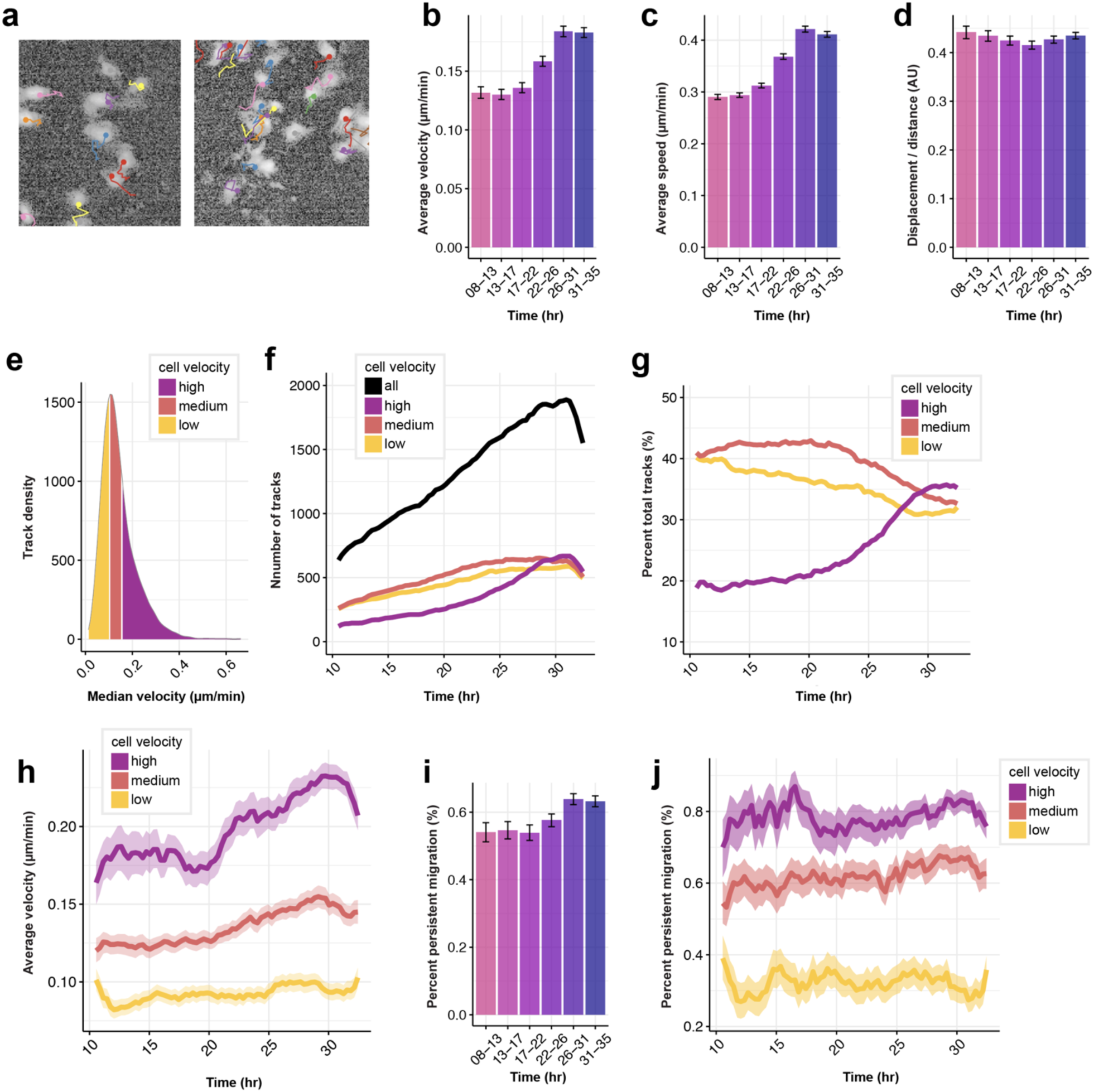
Single-cell tracking of optoWnt cells in optoWnt/WT co-cultures shows increased cell migration and no change in cell persistence. **a)** Representative images of single-cell trajectories of mVenus-NLS+ optoWnt cells. Track color distinguishes different cells in field of view. **b)** Average cell velocities at indicated time intervals after onset of light stimulation. Graph shows mean (>1,000 tracks over 5 fields of view) ± 95% confidence interval. **c)** Average cell speed at indicated time intervals after onset of light stimulation. Graph shows mean (>1,000 tracks over 5 fields of view) ± 95% confidence interval. **d)** Average ratio of cell displacement over distance at indicated time intervals after onset of light stimulation. Graph shows mean (>1,000 tracks over 5 fields of view) ± 95% confidence interval. **e)** Distribution of median cell velocities across all cell tracks. Tracks were binned into three equal groups by median velocity (low: 0-0.15 *µm*/min; medium: 0.15-0.18 *µm*/min; high: 0.18-0.6 *µm*/min) as indicated. **f)** Number of tracks in each median velocity bin over time. **g)** Tracks in each velocity bin over time, shown as percentage of total tracks**. h)** Average cell velocity over time, binned by median cell velocity. Graph shows mean ± 95% confidence interval. **i)** Average percent of cells undergoing persistent migration at indicated time intervals. Graph shows mean percentage ± 95% confidence interval. j) Average percent of cells undergoing persistent migration over time, binned by median cell velocity. Graph shows mean percentage ± 95% confidence interval.

**Extended Data Figure 10.**
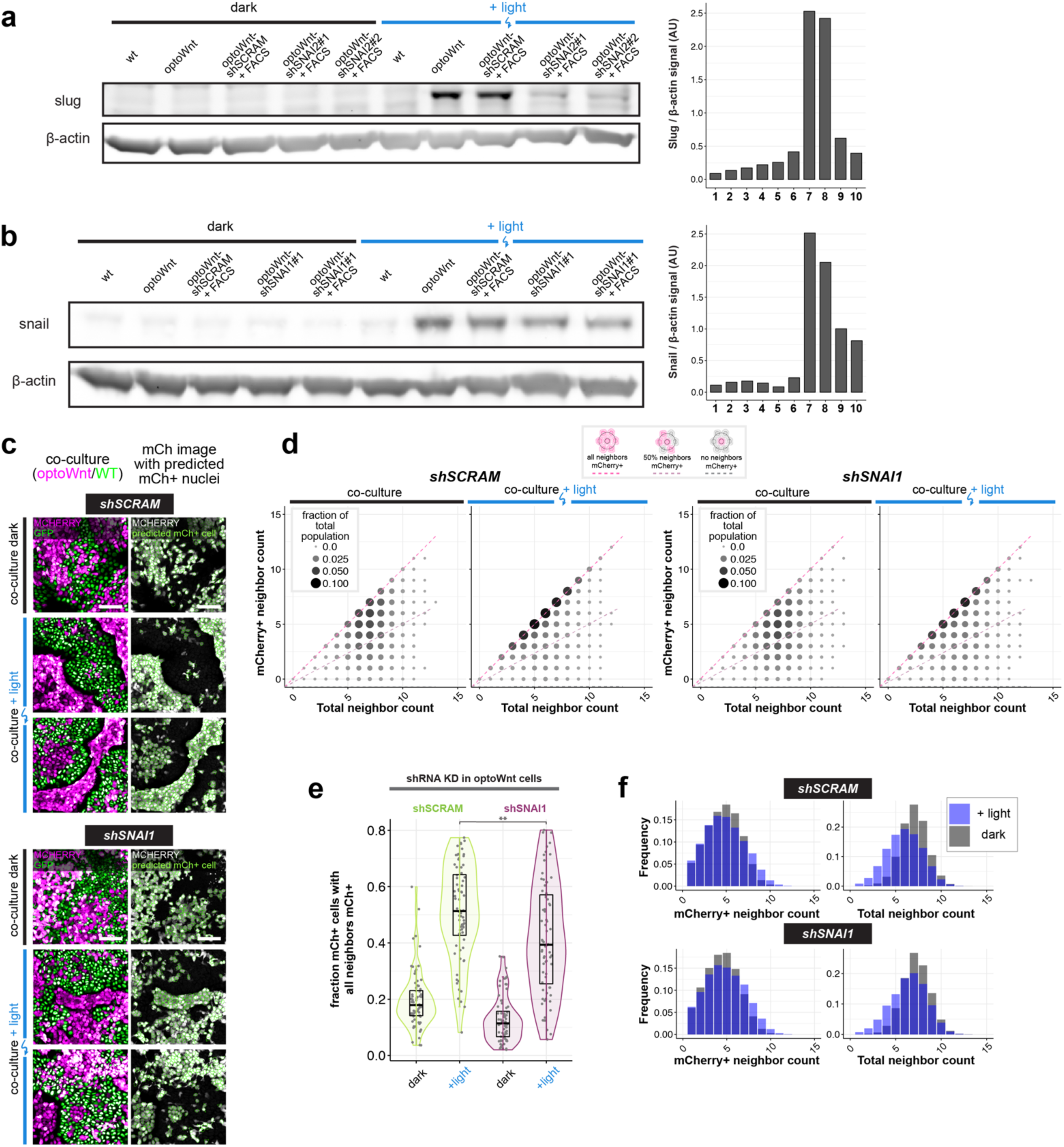
Gene knockdown of EMT regulator SNAI1 in optoWnt cells shows decreased cell self-organization in 2D co-cultures. **a)** Western blot (left) for SLUG protein levels in indicated WT and optoWnt cell lines in response to shRNA knockdown. Cell lines labelled +FACS designates that lines were sorted for GFP+ expression, which marks cells infected with shRNA construct. WB quantification (right), normalized to β-actin loading control. The optoWnt line expressing shRNA snai2 #2 (+FACS) was used for subsequent experiments. **b)** Western blot (left) for SNAIL protein levels in indicated WT and optoWnt cell lines in response to shRNA knockdown. WB quantification (right), normalized to β-actin loading control. shRNA snai1 #1 (+FACS) was used for subsequent experiments. **c)** Representative images of 2D optoWnt/WT co-cultures (left column) in the dark and after 48 hrs illumination. OptoWnt cells express scrambled shRNA (shSCRAM, top panel) or SNAI1 shRNA (shSNAI1, bottom panel). mCherry fluorescence marks optoWnt cells, GFP nuclear fluorescence marks WT cells. Sample images from cell neighbor analysis in CellProfiler (right column) show nuclear outline of mCh+ cells (green) overlaid on mCh channel image. Scale bar 100 *µm*. **d)** Cell neighbor analysis of optoWnt (mCh+) cells in optoWnt-shSCRAM/WT or optoWnt-shSNAI1/WT co-cultures kept the dark or illuminated for 48 hrs. Graph shows the count of total cell neighbors vs. count of mCh+ cell neighbors across total population of analyzed mCh+ cells (72,338 cells analyzed, pooled analysis from n=3 biological replicates). Area and color of points is proportional to the fraction of total population. Constant ratios of mCh+ to total neighbors are highlighted with pink and grey lines. **e)** Quantification of fraction of optoWnt (mCh+) cells whose neighbors are all mCh+ in optoWnt-shSCRAM/WT and optoWnt-shSNAI1/WT co-cultures. Each point represents an analyzed field of view (72 fields of view analyzed, n=3 biological replicates). Unpaired two-samples Wilcoxon test (p = 0.0033). **f)** Histogram of total cell neighbor counts (bottom) and mCh+ cell neighbor counts (top) across all analyzed cells show comparable cell densities between light and dark conditions, as well as between optoWnt-SCRAM/WT and optoWnt-SNAI1/WT co-cultures.

**Table S2.** Primary antibodies used for immunostaining and western blot (WB)

| Antigen | Species | Antibody | Concentration |
| --- | --- | --- | --- |
| LRP6 | Rbt | Ab134146 | 1:1000 1063 |
| BRA | Rbt | Sc20109 | 1:500 1064 |
| β-CATENIN | Rbt | Cell sig 8480 | 1:500 1065 |
| SLUG | Rbt | Cell sig 9585 | 1:2000 (WB) 1066 |
| SNAIL | Ms | Cell sig 3895 | 1:1000 (WB) 1067 |
| β-ACTIN | Ms | Sigma A1978 | 1:5000 (WB) 1068 |
| E-CAD | Gt | RD systems AF748 | 1:500 1069 |
| N-CAD | Rbt | Cell sig 13116 | 1:500 1070 |
| OCT4 | Ms | Sc5279 | 1:500 1071 |
| SOX2 | Rbt | Ab97959 | 1:500 1072 |
| NANOG | Ms | Ab62734 | 1:500 1073 |
| MCHERRY | Ck | Ab205402 | 1:1000 1074 |
| MVENUS | Rbt | Ab6556 | 1:1000 1075 |

**Table S3.**
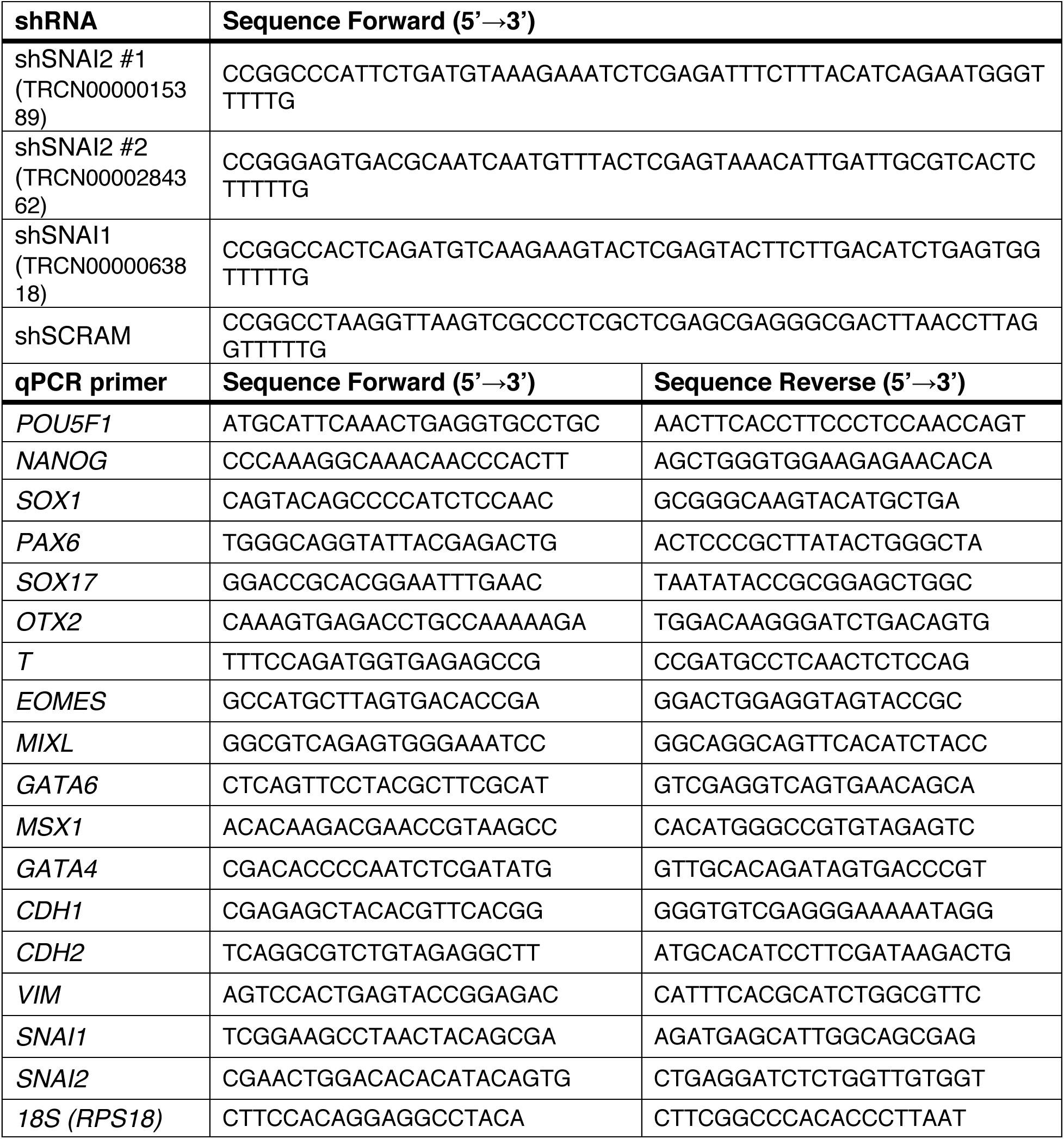
qPCR primers and gene knockdown shRNAs.

## Description of additional supplementary files

File name: TableS1

Description:

**Table S1: Summary of optoWnt hESC RNA sequencing data.** Sample IDs, sequencing depth, and library quality control metrics of RNA-seq study.

File name: SupplementaryVideo1

Description:

**Supplementary Video 1: 3D spheroid timelapse imaging.** Brightfield timelapse imaging of WT, optoWnt, and optoWnt/WT co-culture spheroids under optogenetic stimulation. Time after onset of illumination indicated. Scale bar 100 *µm*.

File name: SupplementaryVideo2

Description:

**Supplementary Video 2: 2D co-culture timelapse imaging.** Timelapse optoWnt/WT co-cultures under optogenetic stimulation (see Fig. 5c). Three image channels displayed: brightfield (BF, grey), CellTracker membrane dye fluorescence (green), and optoWnt-mVenus-NLS cells (magenta). Time after onset of illumination indicated. Scale bar 250 *µm*.

File name: SupplementaryVideo3

Description:

**Supplementary Video 3: 2D co-culture timelapse imaging (zoom).** Zoom-in of two fields of view of Supplementary Video 2. Left: cell aggregation occurs outside hESC colony; right: cell aggregation occurs inside hESC colony. Merge of CellTracker membrane dye fluorescence (green) and optoWnt-mVenus-NLS cells (magenta). Time after onset of illumination indicated. Scale bar 100 *µm*.

File name: SupplementaryVideo4

Description:

**Supplementary Video 4: Aggregate dynamics in 2D co-cultures vs. monocultures.** Brightfield timelapse optoWnt/WT co-culture (left) and optoWnt monoculture (right) under optogenetic stimulation. Magenta arrows highlight aggregates that display collective movement and aggregate fusion. Time after onset of illumination indicated. Scale bar 250 *µm*.

File name: SupplementaryVideo5

Description:

**Supplementary Video 5: Sample single-cell track of optoWnt-mVenus-NLS cell under optogenetic stimulation in optoWnt/WT co-culture.** Timelapse of optoWnt/WT co-culture under optogenetic stimulation, mVenus channel.

File name: SupplementaryVideo6

Description:

**Supplementary Video 6: Multiple single-cell tracks of optoWnt-mVenus-NLS cell under optogenetic stimulation in optoWnt/WT co-culture.** Timelapse of optoWnt/WT co-culture under optogenetic stimulation, mVenus channel.

